# MAPK signaling links BRD2 chromatin occupancy to PI3K/AKT inhibitor sensitivity

**DOI:** 10.64898/2026.09.21.753236

**Authors:** Josefina Mendez, Simone Bruno, Alexandra Indeglia, Tashbib Khan, Isabella L. Ranieri, Jingchao Wang, Jenny Högström, Alissandra L. Hillis, Jonah Lee, John G. Clohessy, Taru Muranen, Wenyi Wei, Franziska Michor, Karen M. Cichowski, Alex Toker

## Abstract

Bromodomain and extra-terminal (BET) proteins, BRD2, BRD3, BRD4, and BRDT, couple histone acetylation to transcription by recruiting elongation and transcription factor complexes to chromatin. Although BET proteins are promising therapeutic targets, the functions of individual family members remain incompletely understood. We identify BRD2 as a co-targetable vulnerability with PI3K/AKT inhibition in breast cancer. Pan-BET inhibition and *BRD2* knockout synergized with PI3K pathway inhibitors in breast cancer cell lines, patient-derived organoids, and *in vivo* models. *BRD2* knockout impaired proliferation of triple-negative breast cancer cells and upregulated signaling and stress-response pathways, including the DNA damage response. Mechanistically, BRD2 is phosphorylated at Ser37 by the mitogen-and stress-activated kinases MSK and RSK, and Ser37 phosphorylation is required for chromatin binding and reader function. Finally, mechanistic digital twin modeling identified BETi–PI3Ki regimens that maintained efficacy while reducing drug exposure. Together, these findings identify BRD2 as a phosphorylation-dependent, co-targetable vulnerability in PI3K-inhibited breast cancer.

## Introduction

Among cancers affecting women, breast cancer is the most frequently diagnosed. Breast cancer is classified into four molecular subtypes based on expression status of the estrogen receptor (ER), progesterone receptor (PR), and human epidermal growth factor receptor 2 (HER2): luminal A, luminal B, HER2-enriched, and triple negative breast cancer (TNBC).^1–5^ TNBC is clinically defined by the absence of ER, PR and HER2 expression, representing approximately 15-20% of all breast cancer diagnoses.^6–10^ Among the four subtypes, TNBC has the worst prognosis, defined by increased metastatic potential and elevated relapse rates.^6–10^ Although select patient populations show benefit with PARP inhibitor therapies or immunotherapy, cytotoxic chemotherapy remains the predominant standard of care.^11–13^ This underscores a pressing clinical need to identify and develop novel targeted therapeutic strategies for this patient population.

Genetic and epigenetic dysregulation of the PI3K/AKT/MTOR pathway is particularly prevalent in breast cancer.^14–18^ Hyperactivation of PI3K/AKT/MTOR promotes tumor cell survival, proliferation, growth and metabolic reprogramming.^18^ In TNBC, gain-of-function alterations in pathway activators, such as PIK3CA (the catalytic subunit of PI3K, p110α; approximately 7% of cases), or loss-of-function alterations in the tumor suppressors PTEN (approximately 35% of cases) and INPP4B (approximately 80% of cases), can occur through mutation, protein loss, or epigenetic silencing.^19–22^ This prevalence has driven efforts to develop pharmacological inhibitors targeting PI3K/AKT.^23, 24^ To date, six PI3K pathway inhibitors have received FDA approval for clinical use. These include the PI3Kα inhibitor GDC-0077 (Inavolisib, Itovebi), approved in combination with the CDK4/6 inhibitor palbociclib and fulvestrant (Faslodex), and the catalytic AKT inhibitor AZD5363 (capivasertib, Truqap), approved in combination with endocrine therapy. Both agents are indicated for hormone receptor positive (HR+), HER2-negative breast cancer patients with PI3K pathway alterations.^25–28^ Despite this progress, the clinical benefit of PI3K/AKT inhibitors as monotherapies across tumor types has been constrained by on-target toxicities and the emergence of acquired resistance.^29–33^ Notably, these agents have also failed to demonstrate efficacy in phase III trials in TNBC when combined with backbone cytotoxic chemotherapy regimens, highlighting the critical need for rationally designed, mechanistically informed therapeutic strategies.

To identify targetable vulnerabilities in TNBC, we previously conducted a genome-wide CRISPR/Cas9 loss-of-function screen under PI3K (BYL719, alpelisib) or AKT (GDC-0068, ipatasertib) inhibition.^34^ Given the established role of chromatin regulation in transcriptional control and cancer cell survival, we specifically examined chromatin regulators within these screening data for genetic dependencies that could enhance sensitivity to PI3K or AKT inhibition. This analysis identified bromodomain-containing protein 2 (BRD2) as a synthetic lethal interactor with both PI3Ki and AKTi. BRD2 belongs to the BET (bromodomain and extra-terminal) domain family which includes BRD3, BRD4 and BRDT. BET family members share a conserved architecture featuring two N-terminal bromodomains (BD1 and BD2), followed by an extra-terminal (ET) domain.^35–37^ Bromodomains are four-helical bundle structures that recognize and bind acetylated lysine (KAc) residues on proteins, including histone proteins. BET proteins act as epigenetic ‘readers,’ facilitating chromatin remodeling and recruiting transcriptional machinery via their ET domains to sites of histone acetylation, thereby regulating gene expression.^38–40^ While BET family members share overlapping functions, they also have distinct, non-redundant roles in transcriptional regulation.^41, 42^ Numerous small molecule BET inhibitors (BETi) have been developed that target the bromodomains of all family members non-selectively, masking potential specific biological functions attributable to individual family members.^37^ Although BRD4 has received the most attention, BRD2 has been implicated in transcriptional initiation, the DNA damage response, and DNA replication.^43–49^ Recently, BRD2 was shown to promote transcriptional initiation via bromodomain-dependent binding to MOF-mediated H4K16ac and C-terminal recruitment of TFIID.^50^ How BRD2 function is regulated in response to upstream signaling, and whether this regulation underlies BRD2-selective dependencies in cancer, remain undefined.

Dysregulation of BET proteins has been implicated in a range of human diseases, including cancer, metabolic and immunological disorders.^51, 52^ Of the pan-BETi that have been developed to date, ZEN-3694 is being investigated in multiple phase 2 trials for solid tumors (NCT03901469, NCT05327010, NCT05607108, NCT04986423). BETi broadly demonstrate pro-apoptotic and anti-proliferative activity across cancer types, including TNBC.^53–63^ One established mechanism underlying the cytotoxicity of BETi is the transcriptional suppression of *MYC,* which is frequently amplified in TNBC.^64^ Although prior studies have reported synergy between PI3K/AKT pathway and BET inhibition in TNBC models,^65, 66^ the mechanistic basis for this synergy and the contribution of BRD2 remain unresolved.

In the present study, we demonstrate that both pan-BET inhibition and BRD2 loss synergize with PI3K and AKT inhibitors in breast cancer cells. We establish BRD2 as a key transcriptional regulator of the stress signaling and the DNA damage response. We further show that BRD2 activity is regulated by growth factor- and stress-stimulated phosphorylation at serine 37 (Ser37) by the mitogen- and stress-activated kinases 1 and 2 (MSK1/2) and p90 ribosomal S6 kinase (RSK). We further integrate longitudinal treatment-response data with mechanistic modeling to optimize BETi–PI3Ki treatment regimens through *in silico* clinical trials. Collectively, these findings position BRD2 as a point of convergence between growth factor signaling and transcriptional control that can be exploited therapeutically in breast cancer.

## Results

### CRISPR/Cas9 screen identifies synergy with combined PI3K/AKT inhibition and *BRD2* knockout

To identify synthetic lethal interactions with PI3K/AKT inhibitors in cancer cells, we previously conducted a genome-wide CRISPR/Cas9 screen in *PIK3CA*-mutant (H1047L) TNBC cells (SUM159).^34^ In this screen, cells were treated for 72 hrs with either vehicle control (DMSO) or cytostatic concentrations of the PI3Kα-selective inhibitor BYL719 (alpelisib, 0.4 µM) or the catalytic AKT inhibitor GDC-0068 (ipatasertib, 3 µM) (**Figure S1A**). Cross-referencing screen hits with the EpiFactors database^67^ revealed 696 epigenetic proteins represented in the screen (**Figure S1B, C**), approximately 320 of which were depleted in each drug treatment arm (**Figure S1D**). These results indicated that a broad range of epigenetic regulators may represent collateral vulnerabilities to either PI3Ki or AKTi. Applying a stringency threshold of log2 fold change ≤ −0.4 narrowed the GDC-0068 arm to 27 epigenetic depleted targets, among which histone readers were disproportionally enriched even after normalizing for class representation in the screen (**Figure S1E**). BRD2, a histone reader belonging to BET protein family, met this threshold in both PI3Ki and AKTi arms of the screen (**Figure S1F**). Notably, BRD2 was the only BET family member to drop out with both anchor drugs, distinguishing it from its paralogs BRD3, BRD4 and BRDT (**Figure 1A**).

**Fig. 1.**
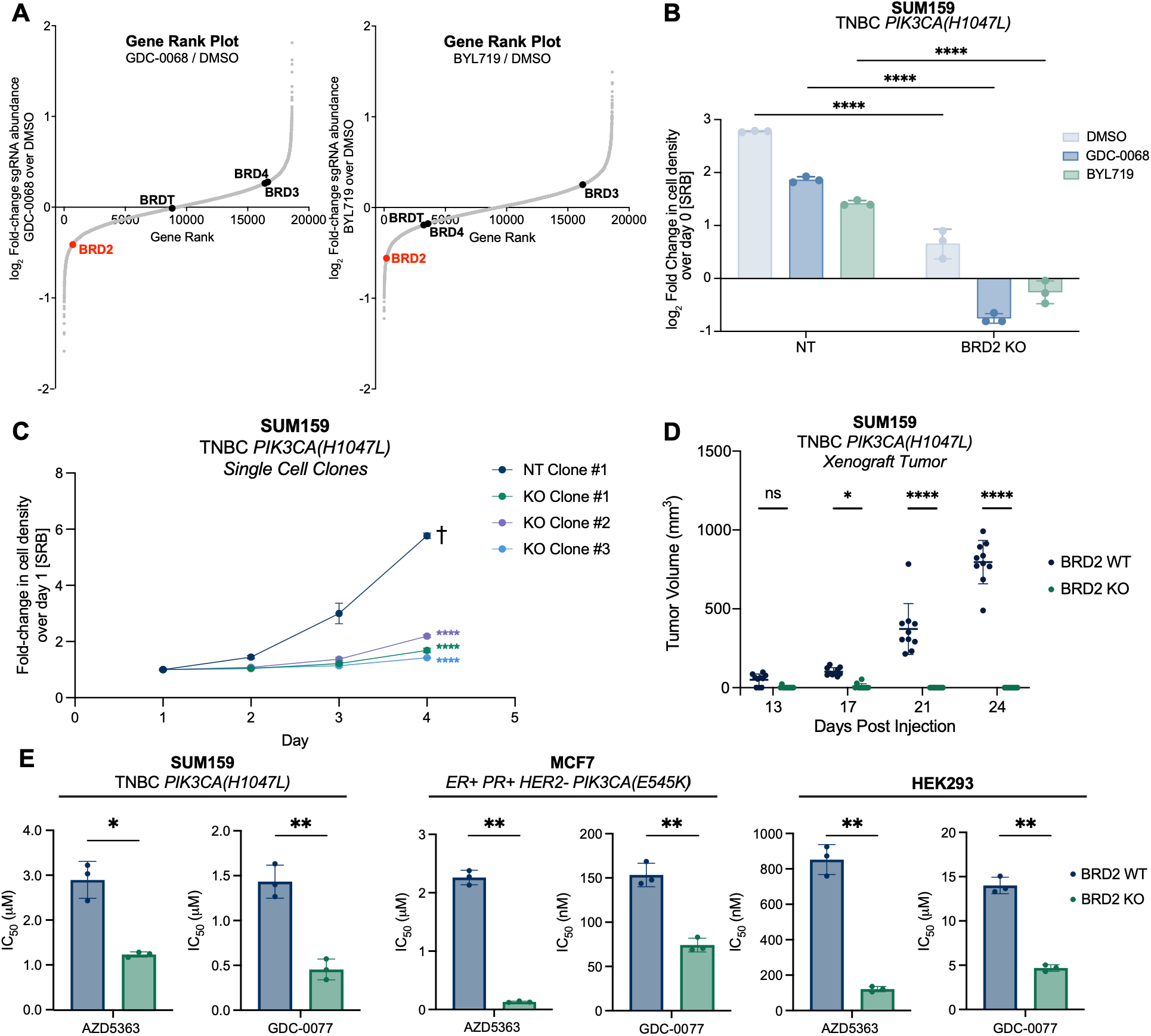
CRISPR/Cas9 screen identifies synergy with combined PI3K/AKT inhibition and BRD2 knockout. (**A**) Rank plots showing the log2 fold-change of each gene plotted against the rank dropout for the BYL719 and GDC-0068 treatment arms compared to the DMSO arm. BET family members are highlighted. (**B**) Non-targeting or *BRD2* knockout (KO) clones were treated with either vehicle or cytostatic doses of the AKTi; GDC-0068 (3 μM) or the PI3Ki; BYL719 (0.4 μM) for 72 hours before harvesting for sulforhodamine B (SRB) assay to determine cell density. Data are represented as mean ± SD (N=3 technical replicates). Statistical analysis was performed using two-way analysis of variance (ANOVA) with Dunnett’s multiple comparison test; asterisks (*) indicate significant differences compared to non-targeting (****, p<0.0001). (**C**) TNBC SUM159 non-targeting (NT) or *BRD2* KO clones were plated, and cell density was measured daily by SRB assay. Data are represented as mean ± SD (N=3 technical replicates). Statistical analysis was performed using two-way analysis of variance (ANOVA) with Dunnett’s multiple comparison test; asterisks (*) indicate significant differences compared to non-targeting on day 4 (****, p < 0.0001). (**D**) SUM159 *BRD2* WT or KO clonal cells were injected orthotopically into the mammary fat pad of NSG mice and tumors were allowed to grow to a palpable size (13 days) before tumor size (mm^3^) was measured every 3-4 days. Data are represented as mean ± SD (N=10 mice). Statistical analysis was performed using two-way analysis of variance (ANOVA) with Dunnett’s multiple comparison test; asterisks (*) indicate significant differences compared to mice with BRD2 WT tumors on day 13, 17, 21 and 24 post-injections. (****, p < 0.0001). (**E**) Dose response analysis of AZD5363 and GDC-0077 in SUM159, MCF7, HEK293 *BRD2* KO and control cells. Data are represented as mean ± SD (N = 3 biological replicates, each comprised of n=3 technical replicates). Statistical analysis was performed using two- tailed Welch’s t-test (**, p = 0.0021).

To validate the screen, we generated single-cell *BRD2* knockout (KO) clones using CRISPR/Cas9 in SUM159 cells (**Figure S1G**). BRD2 loss was cytotoxic when combined with either PI3K or AKT inhibition (**Figure 1B**) and significantly impaired cell proliferation relative to control cells expressing non-targeting gRNA (**Figure 1C**). Similarly, *BRD2* KO significantly suppressed *in vivo* tumor growth in SUM159 xenografts (**Figure 1D**). *BRD2* KO clones were also more sensitive than controls to the clinical AKT and PI3Kα inhibitors capivasertib (AZD5363) and inavolisib (GDC-0077) in SUM159, MCF7, and HEK293 cells (**Figure 1E**). Moreover, suppression of PI3K signaling induced by PI3Ki and AKTi was augmented by BRD2 knockout, evidenced by decreased phosphorylation and total protein levels of the AKT substrate PRAS40 (**Figure S1H**). This suggests that BRD2 loss attenuates the transcription of PI3K pathway genes and their protein products. These results demonstrate that BRD2 loss potentiates the activity of PI3K/AKT inhibitors, at least in part through enhanced suppression of PI3K/AKT signaling.

### BRD2 knockout drives transcriptional reprogramming of proliferation, stress, and DNA repair pathways

To define the transcriptional consequences of BRD2 knockout, we performed bulk RNA-sequencing on wild-type (WT/Non-targeting) and two independent *BRD2* KO SUM159 single-cell clones. Transcriptomic profiles were analyzed using single-sample gene set enrichment analysis (ssGSEA) with MSigDB Hallmark gene sets.^68^ Quality control confirmed strong similarity among the four biological replicates within each condition, while immunoblotting of cell lysates from paired samples verified complete loss of BRD2 protein expression in both knockout clones (**Figure S2A-C**). BRD2 knockout produced widespread transcriptional remodeling, with significant enrichment in 14 of the 50 hallmark gene sets (28%) (**Figure 2A**, **Supplemental Table 1**). Consistent with the established roles of BET proteins in cell-cycle progression and MYC-dependent transcription,^43, 56, 69^ BRD2 loss suppressed E2F targets, MYC targets, and the G2/M checkpoint (**Figure 2A**). A glycolysis gene signature was also significantly reduced following BRD2 knockout, consistent with a recent study implicating BRD2 in the regulation of glycolytic and tricarboxylic acid cycle (TCA) enzymes (**Figure 2A**).^70^ By contrast, gene sets associated with the unfolded protein response (UPR) and apoptosis were significantly upregulated in *BRD2* KO cells (**Figure 2A**). qPCR analysis further validated increased expression of *ATF6* and its downstream target GRP94 (*HSP90B1*), as well as the pro-apoptotic ER stress marker CHOP (*DDIT3*) and Caspase 3. These data demonstrate that BRD2 loss is associated with unresolved ER stress and activation of apoptotic signaling (**Figure 2B**).

**Fig. 2.**
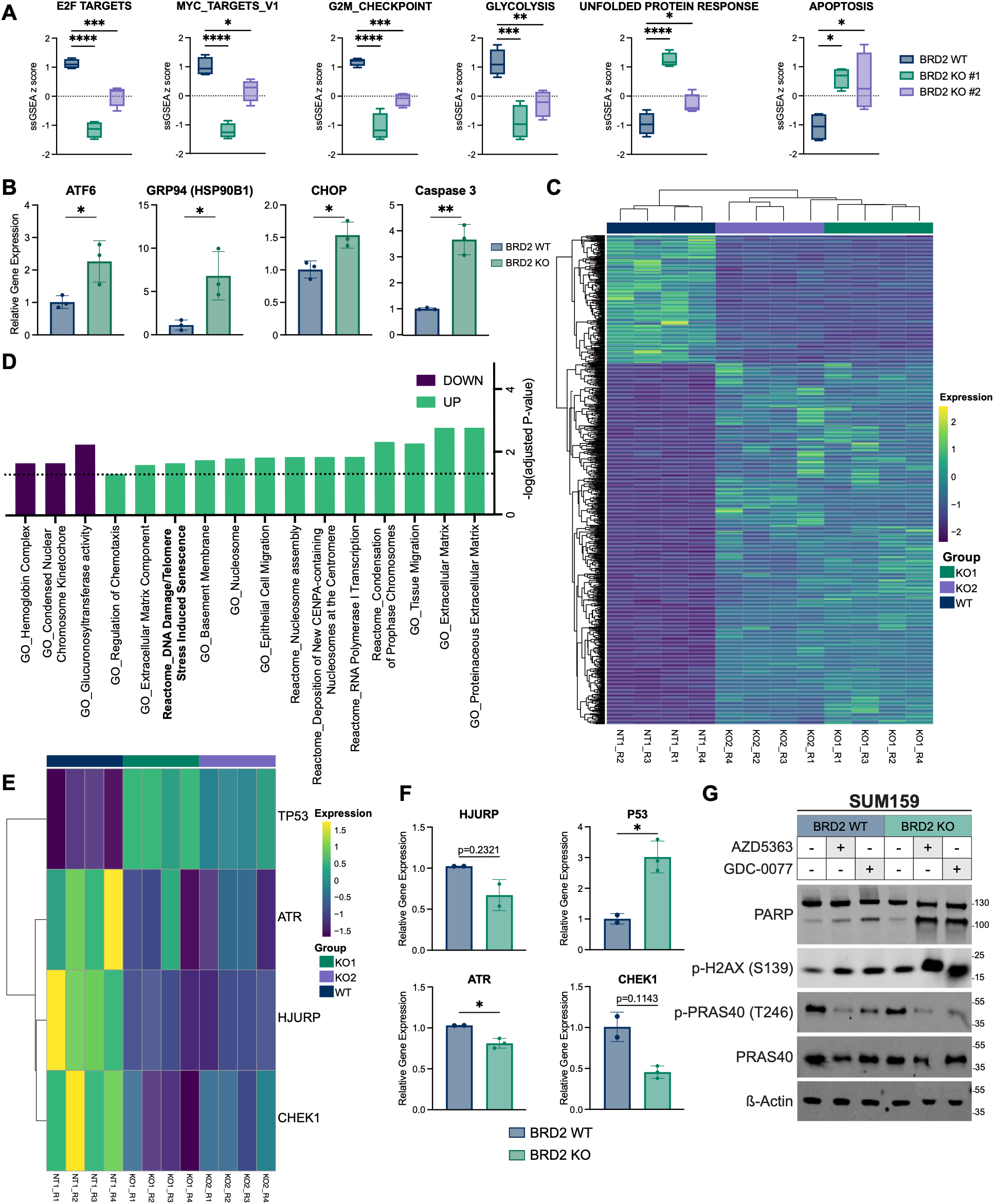
BRD2 knockout drives transcriptional reprogramming of proliferation, stress, and DNA repair pathways. (**A**) ssGSEA z-scores of hallmark gene signatures^68^ in SUM159 single cell *BRD2* WT (non-targeting) or knockout clones (KO1, KO2). Statistical analysis was performed using one-way analysis of variance (ANOVA) with Dunnett’s multiple comparison test (*, p = 0.0332, **, p = 0.0021, ***, p = 0.0002, ****, p < 0.0001). (**B**) Quantitative RT-qPCR analysis of *ATF6*, GRP94 (*HSP90B1*), CHOP (*DDIT3*) and Caspase 3 expression in WT and BRD2 KO (KO Clone 1) SUM159 cells. Gene expression was normalized to 18S housekeeping gene and is presented relative to WT control. Data represent mean ± SD from n=3 technical replicates. Statistical significance was determined using two-tailed Welch’s t-test (*, p = 0.0332, **, p = 0.0021). (**C**) Differential gene analysis identified 822 genes that are differentially expressed between KO1/KO2 and WT cells. Overall results of FPKM cluster analysis are plotted in a heat map clustered using log2(FPKM+1) values. Mainstream hierarchical clustering was implemented to cluster the FPKM values of genes and row Z-scores were homogenized (values generally between −2 and 2). There is only inter-group clustering. (**D**) Gene Ontology (GO) and Reactome pathway analysis^111–113^ of 822 differentially expressed genes between *BRD2* KO (KO1/KO2) and non-targeting. Dotted line indicates cutoff for statistical significance (adjusted p-value < 0.05). (**E**) Overall results of FPKM cluster analysis for WT and KO clones are plotted in a heat map clustered using log2(FPKM+1) values for DNA repair genes. Mainstream hierarchical clustering was implemented to cluster the FPKM values of genes and row Z-scores were homogenized (values generally between −2 and 2). There is only inter-group clustering. (**F**) Quantitative RT-qPCR analysis of TP53, HJURP, ATR, and CHEK1 expression in WT and BRD2 KO SUM159 cells. Gene expression was normalized to 18S housekeeping gene and is presented relative to WT control. Data represent mean ± SD from n=2 or n=3 technical replicates. Statistical significance was determined using two-tailed Welch’s t-test (*, p = 0.0332). (**G**) SUM159 WT and KO cells were treated with cytostatic doses of AZD5363 or GDC-0077 for 72 hours. DNA damage and cell death were assessed by γH2AX and PARP, respectively, and PI3K/AKT inhibition by p-PRAS40.

Differential gene expression analysis performed for each KO clone relative to non-targeting control identified both shared and clone-specific transcriptional changes (**Figure S2D,E**), suggesting a degree of clonal heterogeneity. To prioritize robust BRD2-dependent transcriptional changes and minimize clone-specific effects, subsequent analyses were restricted to genes that were differentially expressed in both knockout clones. This approach identified 822 differentially expressed genes (padj < 0.05, |log_2_FoldChange| > 1), comprising 607 upregulated and 215 downregulated transcripts (**Figure 2C**). Gene Ontology (GO) and Reactome pathway enrichment analyses were performed separately on upregulated and downregulated gene sets to identify biological processes and pathways altered following BRD2 knockout (**Figure 2D**), revealing significant enrichment of genes associated with the DNA damage response and DNA repair. Consistent with this observation, GO and Reactome ssGSEA analysis also revealed significant reduction in gene signatures associated with double stranded break repair and an increase in apoptotic signaling in response to DNA damage in *BRD2* knockout cells (**Figure S2F**). We next evaluated key differentially expressed DNA damage response and checkpoint genes in the RNA-sequencing dataset, revealing reduced expression of Holliday junction recognition protein (*HJURP*), ataxia telangiectasia and Rad3-related protein (*ATR*) and checkpoint kinase 1 (*CHEK1*), together with increased *TP53* transcriptional expression in BRD2 knockouts (**Figure 2E**). qPCR analysis validated these transcriptional changes (**Figure 2F**). Consistent with this transcriptional signature, PI3K/AKT inhibition induced γH2AX more robustly in BRD2 KO cells compared to control, concomitant with enhanced poly(ADP-ribose) polymerase 1 (PARP) cleavage (**Figure 2G**). Together, these findings implicate defective DNA damage surveillance and repair as a basis for the synthetic lethality between BRD2 loss and PI3K pathway inhibition.

### PI3K/AKT inhibition synergizes with BET inhibitors

To determine whether pharmacologic BET inhibition phenocopies the effects of BRD2 knockout, we determined the antiproliferative activity of a panel of clinically relevant BET inhibitors. Across all cell lines tested, BMS-986158 and ABBV-075 exhibited the greatest potency with low nanomolar IC_50_ values (**Figure 3A, Supplemental Table 2**). ZEN-3694 was selected for most subsequent studies because of its advanced clinical development and demonstrated activity in solid tumors.^71, 72^ TNBC (SUM159, HCC70) and HR-positive (MCF7, T47D) breast cancer cells were treated for 72 hrs with cytostatic concentrations of ZEN-3694 and either GDC-0077 or the mutant-selective allosteric PI3Kα inhibitor RLY-2608,^73^ alone or in combination. In all conditions, combined PI3K and BET inhibition produced significantly greater suppression of cell proliferation than either single agent across all cell lines tested. Moreover, combination treatment reduced relative cell density below baseline, suggesting induction of cytotoxic cell death rather than growth arrest alone (**Figure 3B, S3A**). ZEN-3694 dose-dependently sensitized cells to AKT (AZD5363) and PI3Kα (GDC-0077, RLY-2608) inhibition (**Figure 3C**). Highest single-agent analysis scored these interactions as synergistic in two TNBC lines (SUM159, HCC70) and in HR+ lines (MCF7, T47D) (Highest HSA scores: 17.08–48.23; **Figures S3B-J**), indicating that BET/PI3K pathway synergy extends beyond TNBC. Moreover, this synergy extended to two melanoma cell models (Highest HSA scores; SK-MEL-5: 33.82; A-375: 41.7, **Figure S3K-L**).

**Fig. 3.**
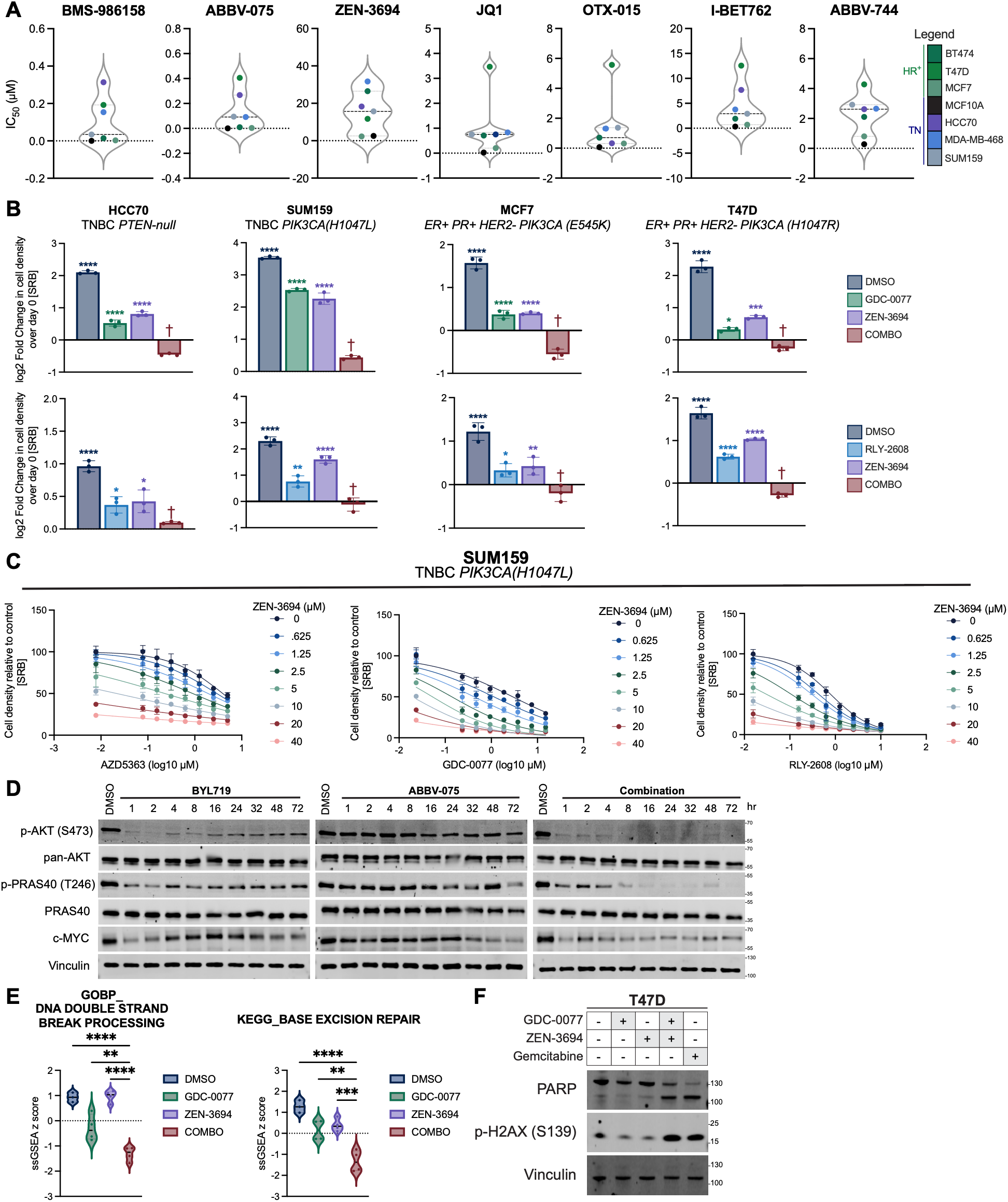
PI3K/AKT inhibition synergizes with BET inhibitors. (**A**) Violin plots depicting the distribution of IC_50_ values (in μM) for individual cell lines treated with a panel of BET inhibitors (JQ1, OTX-015, I- BET762, ABBV-744, ABBV-075, BMS-986158, and ZEN-3694). Each point represents the mean IC_50_ value calculated from three biological replicates (nine technical replicates). (**B**) A panel of TNBC and HR+ breast cancer cell lines were treated with either vehicle or cytostatic doses of the PI3Ki; GDC-0077 (SUM159/HCC70: 1 µM; MCF7/T47D: 15nM), PI3Ki; RLY-2608 (SUM159: 1.25 µM; HCC70: 2.5 µM; MCF7: 1.25 µM; T47D: 156 nM), the BETi; ZEN-3694 (SUM159/HCC70: 5 µM; MCF7/T47D: 2.5 µM), or their combination for 72 hours before harvesting for sulforhodamine B (SRB) assay to determine cell density. Data are represented as mean ± SD (N=3 technical replicates). Statistical analysis was performed using one-way analysis of variance (ANOVA) with Dunnett’s multiple comparison test; asterisks indicate significant differences compared to combo treated cells (†;*, p = 0.0332, **, p = 0.0021, ***, p = 0.0002, ****, p < 0.0001). (**C**) SUM159 TNBC cells were treated with increasing doses of AZD5363 (0-5 µM), GDC-0077 (0-15 µM) or RLY-2608 (0-10 µM) and pan-BETi (BMS-986158: 0-1000 nM, ABBV-075: 0-1000 nM, ZEN-3694: 0-40 µM) for 72 hours, and cell density was measured by SRB assay. Data are represented as mean ± SD (N=3 technical replicates). (**D**) SUM159 cells were treated with the PI3Kα inhibitor (BYL719) (1.5 µM), the pan-BET inhibitor ABBV-075 (500 nM), or their indicated combination. Protein lysates were collected at the indicated time points to assess PI3K pathway signaling. (**E**) ssGSEA z scores of GO_DNA DOUBLE STRAND BREAK PROCESSING and KEGG_BASE EXCISION REPAIR in SUM159 vehicle or drug treated cells (GDC-0077, ZEN-3694 single agent and combination). Statistical analysis was performed using one-way analysis of variance (ANOVA) with Dunnett’s multiple comparison test; asterisks indicate significant differences compared to combo treated cells (*, p = 0.0332, **, p = 0.0021, ***, p = 0.0002, ****, p < 0.0001). (**F**) T47D cells were treated with GDC-0077 (1 µM), ZEN-3694 (10 µM) or their combination. Protein lysates were collected after 72 hours to assess PARP and γH2AX for the induction of cell death and DNA damage, respectively.

To investigate the mechanism underlying this synergy, we examined PI3K pathway signaling following combination treatment. Co-treatment with GDC-0077 and ZEN-3694 further suppressed PI3K pathway activity when compared to PI3Ki alone, with more pronounced reduction in pAKT (Ser473), PRAS40 (pT246) and S6 (pS240/S244) after 48 hrs treatment (**Figure S3M**). Importantly, enhanced pathway suppression was more evident at later time points coinciding with signaling rebound following PI3K inhibitor single agent treatment and was accompanied by enhanced suppression of MYC expression (**Figure 3D**). These findings show that BET inhibition limits adaptive reactivation of PI3K thereby prolonging pathway suppression.

To define the transcriptional changes associated with PI3K and BET inhibition, we performed RNA sequencing following treatment with GDC-0077, ZEN-3694, or the combination. Biological replicates clustered according to treatment, indicating reproducibility with distinct transcriptional responses to each condition (**Figure S3N,O**). Relative to vehicle-treated cells, combination treatment resulted in 5,492 differentially expressed genes (padj < 0.05, |log_2_FoldChange| > 1), including 2,977 upregulated and 2,515 downregulated transcripts (**Figure S3P**). ssGSEA revealed that dual PI3K/AKT BET inhibition uniquely induced pathways associated with the DNA damage response, negative regulation of cell growth, and significantly reduced MYC target gene signatures (**Figure 3E, S3Q**).^74, 75^ We then identified genes whose expression was selectively altered following combination treatment but remained largely unchanged in response to either single agent alone. We identified 204 combination-specific differentially expressed genes, including 66 genes that were downregulated and 138 that were upregulated (**Figure S3R**).

Pathway analysis of these genes revealed a prominent enrichment of DNA-damage-associated processes, including pathways involved in DNA double-strand break repair and base excision repair (**Figure S3R**). Combination treatment was also associated with enrichment of transcriptional programs related to cell-cycle checkpoint activation and cellular senescence, including the G2/M DNA damage checkpoint and DNA damage- and telomere stress-induced senescence (**Figure S3R**). Combination PI3Ki/BETi treatment increased γH2AX, indicative of accumulated DNA damage relative to single-agent treatment (**Figure 3F**). Together, these findings suggest that PI3Ki/BETi imposes a greater genotoxic stress than either treatment alone and may compromise the cellular capacity to appropriately respond to and repair DNA damage. These findings are also consistent with the increased DNA damage observed following genetic BRD2 loss (**Figure 2**), further supporting a role for BRD2 inhibition in promoting DNA damage in the context of PI3K/AKT pathway inhibition.

### PI3K/AKT inhibitors synergize with BET inhibitors to induce cytotoxicity

We next determined whether combined PI3K and BET inhibition promotes cytotoxicity in TNBC (HCC70) and HR+ (T47D) breast cancer cells. Treatment with either GDC-0077 or ZEN-3694 alone produced only a modest increase in propidium iodide (PI)-positivity relative to control. By contrast, combined PI3Ki/BETi resulted in a marked increase in PI-positive cells in both cell models (**Figure 4A**). To determine the kinetics of cell death, we performed IncuCyte longitudinal live-cell imaging. Continuous monitoring of GDC-0077 or ZEN-3694 or combination-treated TNBC cells (HCC70 and SUM159) over five days resulted in a progressive increase in cell death following combination treatment that was significantly greater than either single agent (**Figure 4B**). The combination of PI3Ki and BETi also produced greater suppression of HCC70 xenograft tumor growth than either monotherapy alone, and was well tolerated (**Figure 4C, Figure S4A,B**). We also evaluated the combination in a panel of ER-+ and ER-low primary breast cancer patient-derived organoids (PDOs). Combined PI3K/AKT inhibition (AZD5363/GDC-0077) with ZEN-3694 significantly reduced PDO growth compared with single-agent treatment. This effect was accompanied by decreased EdU incorporation and increased caspase-3 staining, consistent with enhanced apoptotic cell death (**Figure 4D-F, Figure S4C**). Collectively, these data demonstrate that dual PI3K and BET inhibition potently suppresses breast cancer cell growth and drives cytotoxicity.

**Fig. 4.**
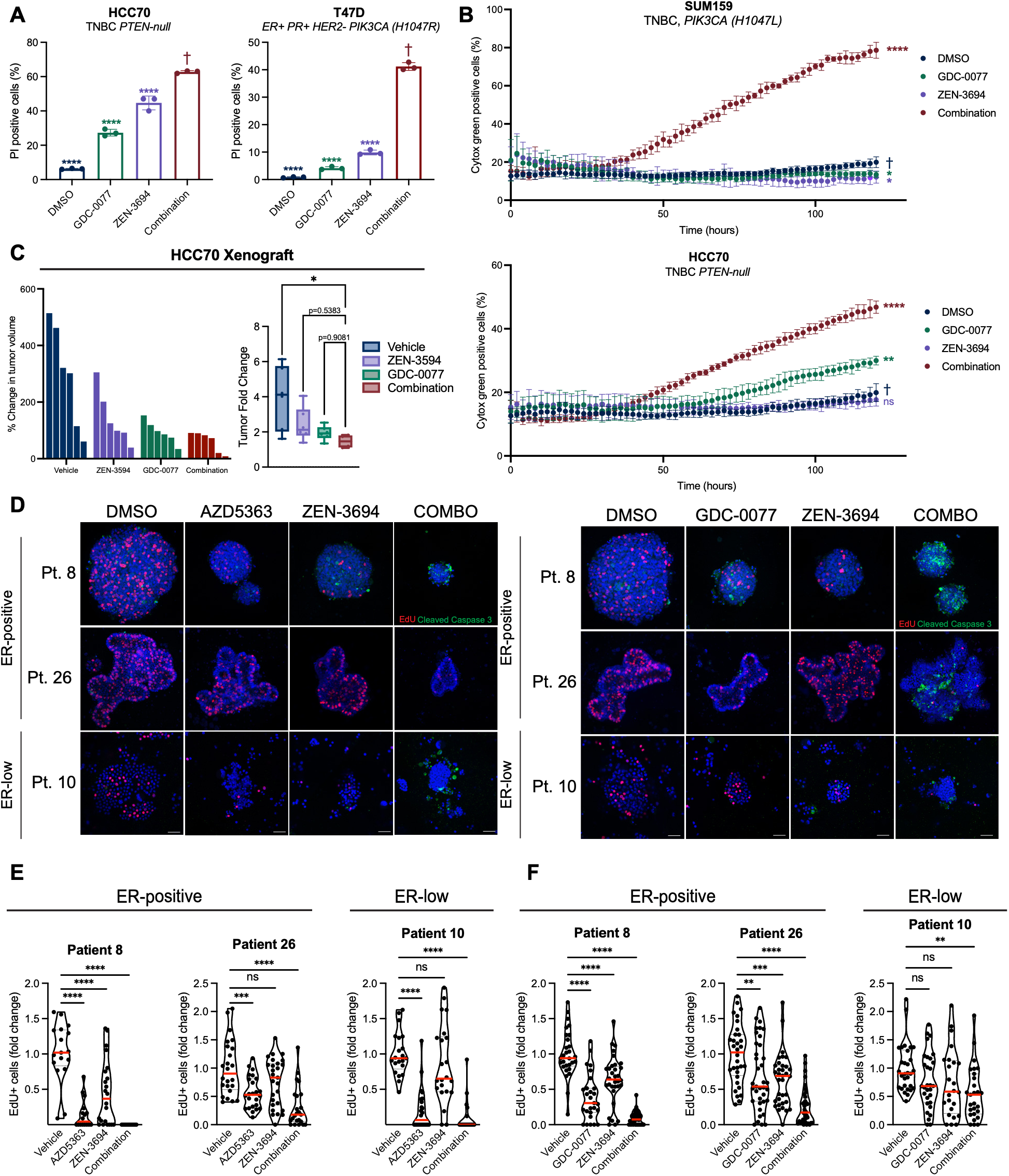
PI3K/AKT inhibitors synergize with BET inhibitors to induce breast cancer cytotoxicity. (**A**) PI death assay showing the percentage of dead cells following treatment with the PI3K inhibitor (GDC- 0077), BET inhibitor (ZEN-3694), or their combination. Triple-negative HCC70 and HR+ T47D cells were treated with GDC-0077 (20 µM and 500 nM respectively), ZEN-3694 (40 µM), or their combination for 72 hours and cell death was assessed by co-staining with Hoechst and propidium iodide (PI) and imaging on a Celigo Imaging Cytometer. Data represent the mean percent cell death ± SD from n=3 biological replicates. Statistical analysis was performed using one-way analysis of variance (ANOVA) with Dunnett’s multiple comparison test; asterisks (*) indicate significant differences compared to combination treated (†) on day 4 (****, p < 0.0001). (**B**) TNBC cell lines (SUM159 and HCC70) were treated with DMSO, GDC- 0077 (SUM159, 10 μM; HCC70, 7.5 μM), ZEN-3694 (SUM159/HCC70, 10 μM), or the combination of GDC-0077 and ZEN-3694. Total cell number (rapid red nuclear dye) and number of dead cells (cytotox green) were measured every 2 hours for 5 days by IncuCyte Live-cell Analysis. Data are represented as mean ± SD of percent cytotox green positive cells (N=4 technical replicates). Statistical analysis was performed using two-way analysis of variance (ANOVA) with Dunnett’s multiple comparison test, asterisks (*) indicate significant differences compared to DMSO (†) at day 5 (*, p = 0.0332; **, p = 0.0021; ****, p < 0.0001). (**C**) Waterfall plot depicting percent change in tumor volume at endpoint (6 days of drug treatment) in SUM159 TNBC xenografts following treatment with vehicle, ZEN-3564 (50 mg/kg, daily), GDC-0077 (25 mg/kg, daily), or their combination. Box-and-whisker plots showing tumor weights at endpoint for each tumor group with each point representing an individual mouse. Statistical analysis was performed using one-way analysis of variance (ANOVA) with Dunnett’s multiple comparison test; asterisks indicate significant differences compared to combo treated cells (****, p < 0.0001). (**D**) A panel of breast cancer PDOs were treated with DMSO, 1 µM AZD5363, 1 µM GDC-0077, 3 µM ZEN-3694 or the combination of GDC-0077/ZEN-3694 or AZD5363/ZEN-3694 for 96 hours and pulsed with EdU and stained with cleaved caspase-3. A representative image for each treatment condition is shown. Scale bar, 50 µm. (**E**) Violin plots showing EdU-positive fold change relative to vehicle for patient derived breast organoids (PDOs) treated with (**E**) DMSO, AZD5363 (1 μM), ZEN-3694 (3 μM), or their combination, or (**F**) DMSO, GDC-0077 (1 μM), ZEN-3694 (3 μM), or their combination. Each data point represents an individual PDO, with median highlighted in red. Statistical analysis was performed using one-way analysis of variance (ANOVA) with Dunnett’s multiple comparison test; asterisks indicate significant differences compared to combo treated cells (*, p = 0.0332, **, p = 0.0021, ***, p = 0.0002, ****, p < 0.0001).

### BRD2 is phosphorylated by MSK and RSK at S37

Post-translational modifications regulate BET protein function, with phosphorylation of BRD4 altering its interactions and transcriptional activity.^76–79^ We therefore asked whether BRD2 is subject to comparable regulation. We interrogated the PhosphoSitePlus^80^ database for conserved kinase phosphorylation motifs within BET family proteins. This analysis identified a highly conserved basophilic-directed AGC kinase consensus motif (RXRXXs/t: R, arginine; X, any amino acid; s/t, phosphorylated serine or threonine)^81^ at position Ser37, a residue previously annotated as phosphorylated in multiple phosphoproteomic studies (**Figure 5A, Figure S5A**). Kinome-wide prediction using the Kinase Library^82^ identified AGC family protein kinases as the highest-confidence candidates for Ser37 phosphorylation (**Supplementary Table 4**). To identify the relevant upstream kinase for Ser37, serum-starved MCF10A cells were stimulated with growth or cellular stress factors in the presence of selective kinase inhibitors (**Figure 5B**). Epidermal growth factor (EGF), phorbol 12-myristate 13-acetate (PMA), and anisomycin induced robust phosphorylation of endogenous BRD2 at Ser37, whereas insulin-like growth factor-1 (IGF1) and insulin did not (**Figure 5C,D, Figure S5B-C**). The kinetics of EGF and anisomycin stimulation of BRD pSer37 were distinct, whereby EGF induced rapid and transient Ser37 phosphorylation, whereas anisomycin produced sustained pSer37 for up to 4 hrs (**Figure 5E, F, Figure S5D,E**), suggesting BRD2 integrates distinct mitogenic and stress-induced signaling outputs.

**Fig. 5.**
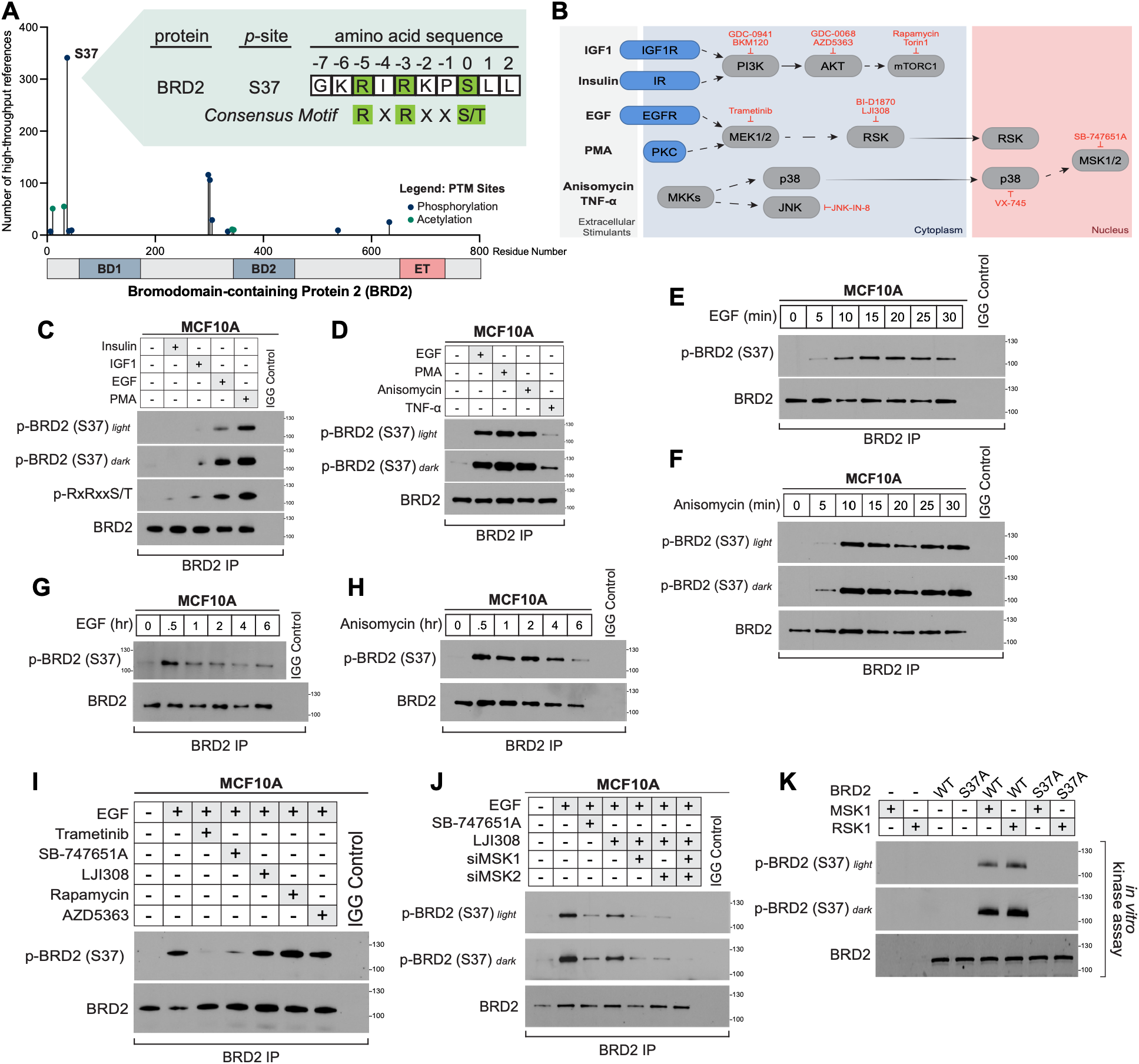
BRD2 is phosphorylated by MSK and RSK at Ser37. (**A**) BRD2 lollipop plot showing functional domains and PhosphoSitePlus-annotated phosphorylation and acetylation sites,^80^ with lollipop height indicating the number of high-throughput references (≥5). (**B**) Diagram of all inhibitors/stimulants used in this study to probe BRD2 phosphorylation. (**C**) BRD2 immunoprecipitation from MCF10A cells following insulin (100 nM), IGF1 (2.6 nM), EGF (10 ng/mL), or PMA (100nM) stimulation (30 minutes). (**D**) BRD2 immunoprecipitation from MCF10A cells following EGF (10 ng/mL), PMA (100nM), Anisomycin (10 µM), and TNF-α (10 ng/mL) stimulation (30 minutes). (**E**) BRD2 immunoprecipitation from MCF10A cells following EGF stimulation (10 ng/mL) for 5–30 minutes. (**F**) BRD2 immunoprecipitation from MCF10A cells following Anisomycin (10 μM) stimulation for 5–30 minutes. (**G**) BRD2 immunoprecipitation from MCF10A cells following EGF stimulation (10 ng/mL) for 30 minutes–6hrs. (**H**) BRD2 immunoprecipitation from MCF10A cells following Anisomycin stimulation (10 μM) for 30 minutes–6hrs. (**I**) BRD2 immunoprecipitation from MCF10A cells following EGF stimulation (10 ng/mL, 30 min) with indicated pathway inhibitors: trametinib (5 µM), SB-747651A (5 µM), LJI308 (10 µM), rapamycin (20 nM), or AZD5363 (5 µM). (**J**) MCF10A cells were transfected with either non-targeting siRNA or siMSK1, siMSK2, or combination siMSK1/2 knockdown for 48 hours before pre-treatment with MSKi and or RSKi and stimulated with EGF (10 ng/mL) for 30 minutes. (**K**) WT or S37A point mutant BRD2 were immunoprecipitated from HEK293 knock-in cells pre-treated with trametinib (5 µM, 30 minutes). IPs were incubated (30 minutes) with purified recombinant full-length MSK1 or RSK1 and ATP at 32 °C.

Although the sequence surrounding Ser37 is consistent with a classical basophilic substrate motif recognized by AGC kinases such as AKT, SGK, and S6K downstream of PI3K, neither insulin nor IGF1 stimulated BRD2 pSer37 (**Figure 5C,D**), and PI3K/AKT/MTOR inhibitors had no effect on EGF-induced phosphorylation (**Figure S5H**). Basophilic-directed kinases including MSK and RSK lie downstream of the MEK/ERK pathway, are stimulated by growth factors including EGF, and are also localized to the nucleus. BRD2 has also been reported to be nuclear,^83^ consistent with its chromatin reader function, and MSK/RSK phosphorylate several chromatin-associated transcriptional regulators.^84–87^ Consistent with this model, EGF-induced pSer37 was sensitive to both MEK inhibition (trametinib) and the MSK inhibitor SB-747651A (**Figure 5I, Figure S5I**). Because SB-747651A inhibits multiple AGC kinases, including RSK family members,^87^ these data suggest that more than one MAPK-activated kinase may contribute to BRD2 phosphorylation. siRNA-mediated depletion of MSK1/2 alone was insufficient to fully suppress BRD2 pSer37 in response to EGF (**Figure S5J**). RSK inhibition alone (LJI308 and BI-D1870) had little effect on BRD2 pSer37 (**Figure S5K**), whereas combined MSK1/2 depletion and RSK inhibition (LJI308) markedly abolished Ser37 phosphorylation (**Figure 5J, Figure S5L,M**). Finally, both MSK1 and RSK1 directly phosphorylated BRD2 in an *in vitro* protein kinase assay (**Figure 5K**). These data support a mechanism by which MSK1/2 and RSK function redundantly to phosphorylate BRD2 at Ser 37.

We also determined whether p38-MSK stress signaling engages a similar regulatory mechanism. Anisomycin-induced BRD2 Ser37 phosphorylation was insensitive to RSK (LJI308) or JNK (JNK-IN-8) inhibition and was selectively blocked by p38α inhibition (VX-745) (**Figure S5O,P**). Together, these data show that BRD2 Ser37 functions as a convergent substrate of MAPK signaling, integrating mitogenic ERK signaling through redundant MSK/RSK activity, and stress-induced p38 signaling primarily through MSK. Regulation of BRD2 pSer37 downstream of MEK and MSK/RSK was conserved across a multiple breast cancer and melanoma cell lines (**Figure S5Q-V**). Moreover, combined inhibition of MSK and RSK (SB-747651A + LJI308) phenocopied *BRD2* loss and BETi by synergizing with PI3Ki (GDC-0077) and AKTi (AZD5363) (**Figure S5W,X**). Collectively, these findings identify BRD2 Ser37 as an MSK/RSK substrate, supporting a model in which MAPK-driven signaling converges on BRD2 to sustain adaptive transcriptional programs under PI3K pathway suppression.

### Phosphorylation of BRD2 at Ser37 regulates its chromatin occupancy

To determine the functional role of BRD2 Ser37 phosphorylation, we generated HEK293 knock-in (KI) cell lines expressing phospho-deficient BRD2 S37A or phosphomimic S37D. Immunofluorescence analysis showed that BRD2 was constitutively nuclear in all genotypes and Ser37 mutation to either Ala or Asp did not alter its subcellular localization (**Figure S6A**). We next asked whether Ser37 phosphorylation instead regulates BRD2 chromatin binding. We performed BRD2 ChIP-seq in HEK293 KI cell lines expressing BRD2 WT, S37A, S37D, with *BRD2* KO as control, along with IgG and H3K27ac ChIP-seq in the WT background as additional controls. Principal component analysis (PCA) of peak signal showed that WT and S37D replicates clustered together, indicating that the phosphomimetic mutant more closely matches the WT chromatin-binding profile compared to the S37A phospho-deficient BRD2 mutant (**Figure S6B**). Global BRD2 chromatin occupancy was markedly altered by Ser37 mutation, whereby BRD2 S37A bound approximately half as many genomic sites as WT. By contrast, BRD2 S37D bound more sites than WT, including a subset of new peaks not evident in WT BRD2 (**Figure 6A,B**). Ser37 phosphorylation is therefore required to maintain the full breadth of BRD2 occupancy across the genome.

**Fig. 6.**
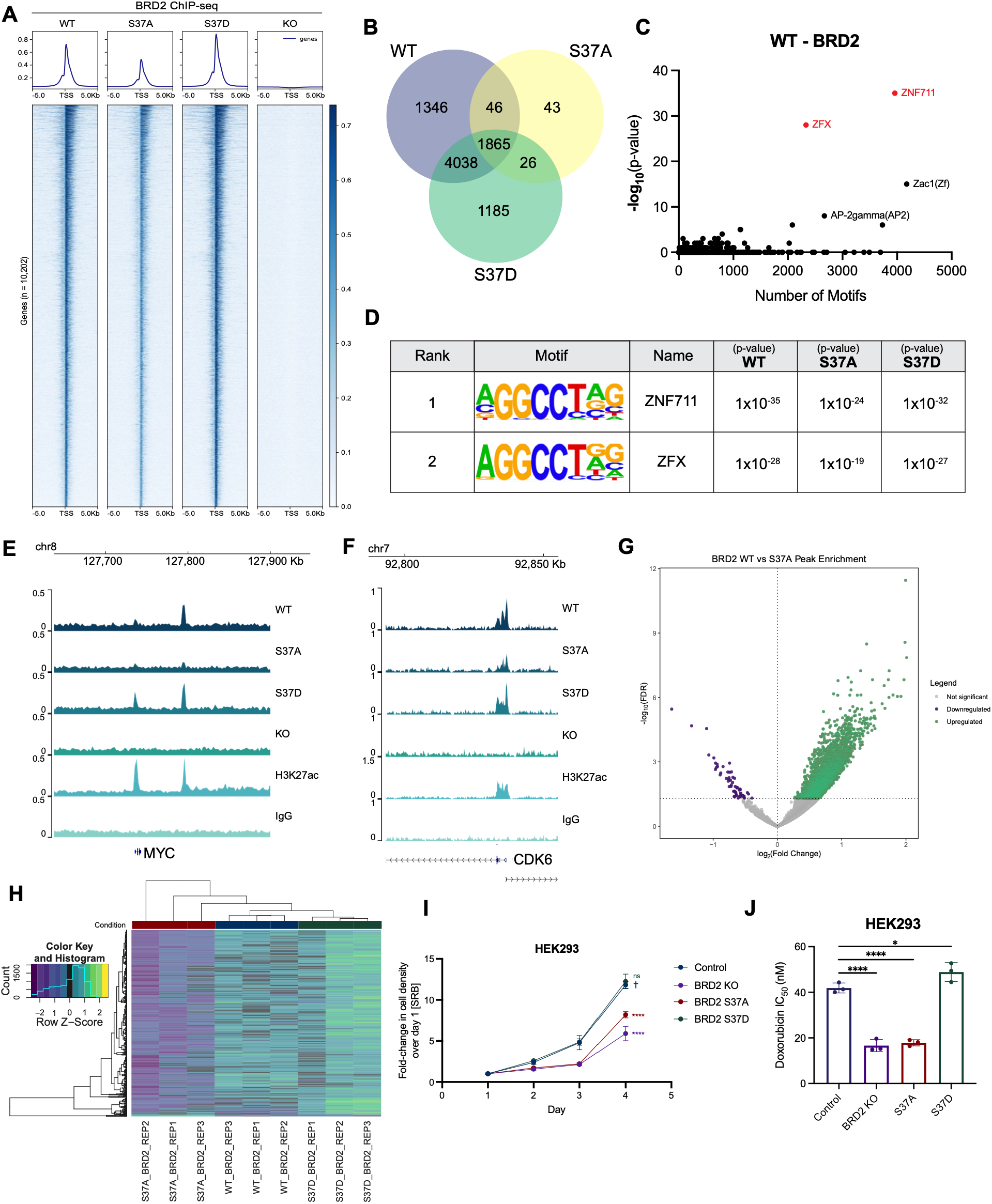
Phosphorylation of BRD2 at Ser37 regulates its chromatin occupancy. (**A**) Heatmaps of BRD2 binding (ChIP-seq) in HEK293 WT, *BRD2* KO, BRD2 S37A, and BRD2 S37D knock-in cells. BRD2 binding is shown for n = 10,202 genes. Data are representative of n = 3 independent replicates. Average density profiles of BRD2 binding are also plotted above the heatmaps. (**B**) Venn diagram of overlapping and unique BRD2 peaks identified in BRD2 WT, S37A, and S37D cells. (**C**) HOMER motif analysis of BRD2 peaks in HEK293 WT cells. (**D**) The top 2 motifs from motif analysis of BRD2 WT, S37A, S37D peaks using HOMER show ZNF711 and ZFX as the most significantly enriched TF motifs at BRD2 sites. p-values for each cell line are shown. (**E**) ChIP-seq genome browser screenshot of BRD2 binding in HEK293 WT, S37A, S37D, KO cells at the *MYC* gene. (**F**) ChIP-seq genome browser screenshot of BRD2 binding in HEK293 WT, S37A, S37D, KO cells at the *CDK6* gene. (**G**) Volcano plot of differential binding analysis of BRD2 WT and S37A peak enrichment. Significant upregulated peaks (n = 4604) are labeled in green, significant downregulated (n = 59) in purple, and non-significant in gray. (**H**) Heatmap clustering analysis of top 1000 genes that show differential binding across BRD2 WT, S37A, and S37D cells. (**I**) HEK293 WT, *BRD2* KO, or BRD2 S37A/S37D clones were plated, and cell density was measured daily by SRB assay. Data are represented as mean ± SD (N=3 technical replicates). Statistical analysis was performed using two-way analysis of variance (ANOVA) with Dunnett’s multiple comparison test; asterisks (*) indicate significant differences compared to *BRD2* KO on day 4 (****, p < 0.0001). (**J**) Dose response analysis of doxorubicin in HEK293 *BRD2* WT, KO, S37A and S37D cells. Data are represented as mean ± SD (N = 3 biological replicates).

Across all genotypes, the majority of BRD2 peaks localized to gene promoters, and this distribution was unaffected by S37A or S37D mutation (**Figure S6C**). Hypergeometric Optimization of Motif EnRichment (HOMER) motif analysis identified the paralogous C2H2 zinc-finger proteins ZNF711 and ZFX as the top enriched motifs, with consistent enrichment across all three replicates for WT, S37A and S37D cells (**Figure 6C,D, Figure S6D**). These factors mark highly active, nucleosome-depleted CpG-island promoters that regulate proliferation, consistent with the established growth-promoting function of BRD2.^88, 89^ At representative proliferation-associated genes *MYC* and *CDK6*, BRD2 peak intensity was reduced in S37A cells relative to WT, while S37D showed a modest increase (**Figure 6E,F**). This corresponded with reduced *MYC* and *CDK6* transcript levels in S37A cells by qPCR, while WT and S37D levels were comparable (**Figure S6E**). We used DiffBind with DESeq2 to identify regions of differential BRD2 binding across genotypes. Consistent with global peak counts, most sites differing between WT and S37A reflected reduced binding in S37A, whereas WT and S37D differed at comparatively few sites (**Figure 6G, Figure S6F**). Unsupervised clustering grouped S37D with WT and separated both from S37A, indicating that the phosphomimetic BRD2 mutant largely retains wild-type binding behavior. (**Figure 6H**). Ser37 phosphorylation therefore serves to regulate the extent of BRD2 chromatin occupancy without altering the genomic features of sequence motifs of BRD2-bound regions.

Finally, given the magnitude of BRD2 chromatin redistribution observed with mutation of Ser37, we determined the functional consequences for BRD2-mediated cell proliferation. BRD2 S37A cells exhibited significantly impaired proliferation, comparable to *BRD2* KO. By contrast, S37D cell proliferation was indistinguishable from WT BRD2 (**Figure 6I**). Because BRD2 regulates both the DNA damage response and cellular metabolism (**Figure 6A**), we asked whether either process underlies the S37A growth defect. Doxorubicin IC_50_ was reduced 2-fold in *BRD2* KO and in S37A cells relative to WT, while S37D cells were unchanged (**Figure 6J**). These results indicated that Ser37 phosphorylation contributes to the BRD2-mediated DNA-damage response. BRD2 has been reported to regulate glycolytic and TCA cycle enzymes. ^70^ Consistent with this, RNA-seq of *BRD2* KO cells revealed coordinate downregulation of glycolysis gene sets (**Figure 2A**). Metabolomic profiling showed a corresponding reduction in glycolytic and TCA cycle intermediates in both *BRD2* KO and S37A cells, which was largely restored by S37D (**Figure S6G**). Specifically, both *BRD2* KO and S37A showed a marked depletion of glycolytic intermediates, including phosphoenolpyruvate, 3-phosphoglycerate, and fructose-1,6-bisphosphate, and of TCA cycle intermediates including isocitrate and malate (**Figure S6G**). Taken together, these results demonstrate that Ser37 phosphorylation maintains BRD2-dependent DNA damage responses and metabolic homeostasis, and that loss of these functions may contribute to the proliferation defect seen with *BRD2* KO and S37A mutation.

### A longitudinal digital twin model enables prediction and optimization of BETi–PI3Ki treatment

Having established the therapeutic interaction between BET and PI3K inhibition and investigated its underlying mechanisms, we next used these findings to investigate the predicted performance of this combination and identify optimized treatment regimens using a computational modeling platform. To this end, we integrated the longitudinal treatment-response imaging data (**Figure 4B** and **Figure S7A,B**) with a mechanistic digital twin framework previously developed.^90^ The resulting calibrated model was coupled with human pharmacokinetic models to conduct *in silico* clinical trials across candidate ZEN-3694 and GDC-0077 treatment schedules (**Figure 7A**).

**Fig. 7.**
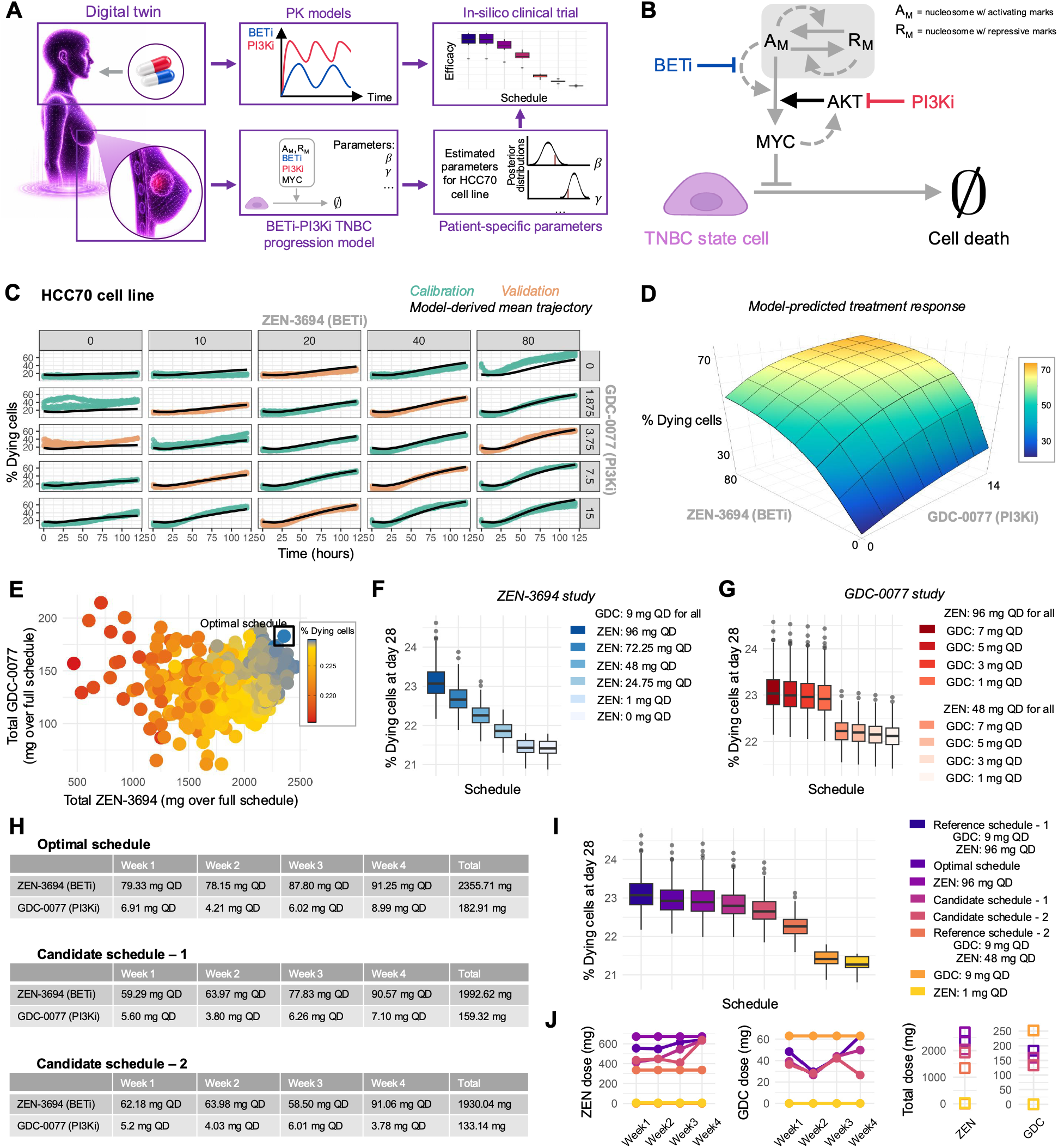
Optimization of BETi–PI3Ki regimens through *in silico* clinical trials using a digital twin model. (**A**) Overview of the digital twin and *in silico* clinical trial framework. (**B**) Schematic diagram of the BETi-PI3Ki TNBC progression model, describing TNBC cell death and its regulation by MYC, chromatin modifications, BETi and PI3Ki. (**C**) Model calibration and validation using longitudinal measurements of treatment-induced cell death across BETi-PI3Ki conditions. Green and orange denote experimental data used for model calibration and validation, respectively, and black denotes the posterior mean model prediction. Drug concentrations are reported in µM. (**D**) Model-predicted response surface across ZEN- 3694 (x-axis) and GDC-0077 (y-axis) concentrations. Treatment efficacy is quantified as the percentage of dying cells at 120 hours (z-axis). (**E**) Candidate treatment schedules (*N* = 2068) evaluated in the *in silico* clinical trial using a genetic algorithm, shown according to total ZEN-3694 and GDC-0077 exposure. The black square indicates the optimized regimen. (**F,G**) Treatment efficacy, expressed as the percentage of dying cells at day 28 (end of one treatment cycle), across schedules with a fixed GDC- 0077 dose and varying ZEN-3694 doses (**F**), or a fixed ZEN-3694 dose and varying GDC-0077 doses (**G**). (**H**) Summary of the optimal and candidate schedules considered in panels I and L. (**I**) Treatment efficacy for the reference schedules, the optimal schedule, two additional candidate schedules, and single-drug schedules. For panels (**F**), (**G**), and (**I**), boxes represent the interquartile range, with the median indicated by the horizontal line. Whiskers extend to the minimum and maximum values excluding outliers, shown as dots. Each regimen is represented with a different color and evaluated across 500 *in silico* patients. (**J**) Total drug exposure for each regimen considered in panel I. For both drugs, dots represent the total dose delivered during each treatment week, and squares represent the overall cumulative dose across the entire treatment cycle.

To describe the response of TNBC cells to BETi and PI3Ki, we developed a mechanistic BETi– PI3Ki TNBC progression model (**Figure 7B**, **STAR Methods, and Eq. (1)**), in which viable cells transition to cell death under the regulation of MYC. In this model, MYC expression is controlled by a chromatin modification circuit, adapted from previous studies,^91–93^ describing the competing establishment and maintenance of activating and repressive histone modifications. BET proteins promote MYC transcription through its interaction with active chromatin, while BET inhibition suppresses this process.^37, 56^ The model also incorporates the experimentally observed attenuation of PI3K pathway reactivation by BET inhibition following PI3K inhibition (**Figure 3D**).

We next calibrated and validated the model using longitudinal live-cell imaging data from HCC70 cells (**Figure 7C**). Model parameters were estimated from the training data using Bayesian inference.^90, 94^ The model accurately reproduced the training data (green; Pearson *r* = 0.873, normalized RMSE = 2.8) and maintained comparable accuracy on treatment conditions excluded from parameter estimation (orange; Pearson *r* = 0.931, normalized RMSE = 2.4), demonstrating its predictive capability. The calibrated model was then used to characterize treatment efficacy across BETi and PI3Ki combinations (**Figure 7D**). GDC-0077 alone induced limited cell death, whereas increasing ZEN-3694 concentrations produced greater efficacy, consistent with the experimental data (**Figure 4A**). Combining the inhibitors further enhanced efficacy, with the greatest response at high concentrations of both drugs.

We then conducted an *in silico* clinical trial to identify combination schedules that maximize efficacy while limiting drug exposure (**Figure 7E).** Patient heterogeneity was captured through pharmacodynamic parameter variability^90^ and ZEN-3694 and GDC-0077 pharmacokinetics were modeled using available data (**STAR Methods**). Candidate 28-day schedules were optimized using a genetic algorithm,^95^ allowing the dose of each drug to vary weekly within clinically relevant ranges (**Figure S7C**) For comparison, we considered reference regimens combining GDC-0077 at 9 mg QD with ZEN-3694 at either 48 or 96 mg QD, based on clinically investigated doses and tolerability.^96, 97^ Analysis of the simulated schedules showed increasing efficacy with increasing doses of either ZEN-3694 or GDC-0077, with a greater sensitivity to the ZEN-3694 dose (**Figure 7F,G**). The *in silico* clinical trial identified an optimized regimen with efficacy comparable to the high-dose reference, but with lower overall ZEN-3694 and GDC-0077 exposure achieved by varying the doses across treatment weeks (**Figure 7H-J**). Similar results were obtained from *in silico* clinical trials in a heterogeneous virtual population obtained by incorporating heterogeneous close based on data from both HCC70 and SUM159 cells (SI-Figure S10).

## Discussion

In this study, we identified BRD2 in an unbiased genome-wide CRISPR screen as synthetically lethal with both PI3K and AKT inhibition in TNBC. BRD2 was the only BET family member depleted under both conditions, indicating a non-redundant role for BRD2 in maintaining cell viability during PI3K pathway suppression. This selectivity is notable given the sequence similarity among BET bromodomains and the long-standing assumption of functional redundancy within the family. Further, we demonstrate that BRD2 chromatin occupancy is governed by mitogen- and stress-activated kinase signaling through phosphorylation at Ser37, a residue found in BRD2 but not in BRD3, BRD4, or BRDT. This places BRD2 at the interface between MAPK-driven signaling and the transcriptional programs that sustain viability under PI3K pathway suppression and identifies a regulatory feature that distinguishes BRD2 from paralogs whose bromodomains are otherwise engaged indiscriminately by BET inhibitors. Oncogenic kinase signaling is known to converge on chromatin, but characterized examples largely involve writers, erasers, or remodelers. For example, AKT phosphorylates EZH2 to suppress H3K27 trimethylation,^98^ and KMT2D to suppress H3K4 mono- and di-methylation and thereby ER-dependent transcription.^99^ These results parallel our findings, as PI3Kα inhibition relieves KMT2D phosphorylation and releases a compensatory ER transcriptional program that limits drug efficacy. Further, canonical signaling pathways have been known to act directly on histones, such as MSK/RSK phosphorylation of histone H3 at Ser10.^100–102^ Direct kinase regulation of a BET reader, by contrast, has been described almost exclusively for BRD4, which has been characterized as a substrate of several kinases including CK2, JAK2 and CDK1.^76–79^ Our data instead place BRD2 downstream of ERK and p38 via MSK and RSK, identifying a mitogen-responsive kinase cascade that regulates the genome-wide chromatin occupancy of an acetyl-lysine reader. BRD2 is therefore better understood not as a constitutive chromatin scaffold, but as a dynamic, signal-responsive chromatin reader.

Importantly, the fact that both MSK and RSK phosphorylate Ser37 is consistent with the substrate promiscuity of the AGC kinase family. MSK1/2 and RSK1-4 share a basophilic kinase motif and converge on a common set of nuclear substrates including as RelaA/p65 NF-κB^85^ and ER8.^86, 87^ This redundancy predicts that inhibition of a single kinase is insufficient to abolish Ser37 phosphorylation, consistent with our reliance on the multi-AGC inhibitor SB-747651A, and implies that the dominant kinase may differ between growth factor-and stress-driven contexts. This is particularly relevant under PI3K pathway inhibition, where relief of negative feedback on receptor tyrosine kinases produces well-documented ERK reactivation.^30, 103, 104^ Whether Ser37 phosphorylation is sustained or increased during ERK rebound, and therefore whether BRD2 is an active participant in adaptive resistance rather than a passive requirement, remains an open question. Further, Ser37 phosphorylation could in principle regulate BRD2 function through several distinct mechanisms, including alter subcellular localization or regulate the localization of BRD2 to distinct genomic loci. In fact, several phosphorylation sites on BRD4 have been characterized to regulate its chromatin binding, recruitment of partner transcription factors, or localization to certain gene loci.^76–79^ However, our data showed that phosphorylation at BRD2 S37 does not alter BRD2 nuclear localization or overall genomic binding preference, but instead regulates the breadth of BRD2 chromatin occupancy. A change in acetyl-mark recognition capabilities would be expected to redistribute BRD2 to a different set of loci, whereas a reduction in binding strength would preferentially deplete BRD2 from weakly occupied sites while preserving the strongest, which is what we observe. We therefore favor a model in which Ser37 phosphorylation tunes the avidity of the tandem bromodomains, and thus chromatin residence time, rather than which acetylation marks BRD2 reads. Testing this will require direct kinetic measurement of BRD2 chromatin binding, quantitative binding assays comparing phosphorylated and unphosphorylated BRD2 against defined acetylated nucleosomes, and structural or HDX-MS analysis of the phosphorylation-dependent conformational change. Importantly, this regulatory mechanism translated into functional consequences, whereby loss of Ser37 phosphorylation phenocopied complete BRD2 loss, impairing cell proliferation, increasing sensitivity to DNA-damaging agents, and depleting key anabolic metabolites. Together, these findings position BRD2 as a signaling-responsive epigenetic regulator, integrating extracellular and intracellular cues to sustain cancer cell proliferation and maintain cell state.

Our mechanistic findings also point to a viable combination strategy for treatment. Pan-BET inhibitors largely phenocopied BRD2 loss, cooperatively suppressed DNA repair and more effectively inhibited tumor growth in TNBC xenografts and in ER+ and ER-low patient-derived breast organoids. While the mechanisms underlying this synergy are likely multifaceted, we found that combination treatment enhanced suppression of PI3K pathway signaling and prevented the adaptive signaling rebound seen with PI3K inhibitor monotherapy. These findings indicate that BET-dependent transcription supports adaptive resistance to PI3K pathway inhibition, and that co-targeting this dependency may deepen and prolong therapeutic responses. While pan-BET inhibitors have shown clinical activity across tumor types, they are limited by a narrow therapeutic index, with on-target gastrointestinal toxicity and thrombocytopenia limiting sustained exposure.^105^ However, ZEN-3694 is under active investigation in breast cancer, including in combination with talazoparib and with pembrolizumab plus nab-paclitaxel in metastatic TNBC (NCT03901469, NCT05422794). Our findings suggest two routes to a wider therapeutic window. The direct route is BRD2-selective inhibition or degradation, which our data predict would retain the synergy with PI3K/AKT inhibition while sparing BRD4-dependent functions in normal tissue. A more indirect route is upstream kinase inhibition, and because Ser37 has no counterpart in BRD3, BRD4, or BRDT, this route confers a paralog selectivity that bromodomain-directed inhibitors cannot achieve. Attenuating MSK and RSK activity would be expected to reduce BRD2 chromatin occupancy while leaving other BET family functions intact. Indeed, we demonstrate that that pharmacologic inhibition of MSK and RSK robustly synergizes with PI3K and AKT inhibitors, potentially offer a therapeutic alternative with a distinct toxicity profile to BET inhibitors. Notably, ZEN-3694 is already being evaluated in combination with the MEK inhibitor binimetinib in RAS-pathway-altered solid tumors and TNBC (NCT05111561). Our results provide a mechanistic rationale for that combination, since MEK inhibition would reduce ERK-dependent MSK and RSK activity and consequently BRD2 Ser37 phosphorylation. Direct RSK inhibition is also clinically tractable: PMD-026, a first-in-class oral pan-RSK inhibitor, is in Phase 2 evaluation in metastatic breast cancer with RSK2 immunohistochemistry used for patient selection (NCT04115306). Because the PI3K/AKT partner agents are now standard of care in breast cancer following the approvals of capivasertib and inavolisib,^25–28^ the combinations we describe are testable in trials.

Our data suggest impaired DNA damage repair contributes to the synthetic lethality between BRD2 loss and PI3K pathway suppression. Specifically, knockout of BRD2 reduced the levels of ATR and CHEK1 (Chk1), two central mediators of the replication stress checkpoint,^106^suggesting that BRD2 loss may impair mechanisms that maintain genome stability during DNA replication. These findings align with prior work showing that BRD2 facilitates resolution of R-loop-associated DNA damage and replication stress.^107, 108^ However, our data also implicate BRD2 loss in the disruption of several additional survival programs, including MYC-dependent transcription, anabolic metabolism, and activation of the UPR and other cell stress pathways. Collectively, these findings support a model in which BRD2 loss disrupts DNA damage repair alongside broader survival and stress-response programs, rendering cells particularly vulnerable to PI3K pathway suppression.

Finally, having established and mechanistically characterized the therapeutic interaction between BET and PI3K inhibition, we then used a digital twin model to investigate the efficacy of different combination treatment schedules and to identify best therapeutic strategies. These analyses identified a candidate BETi–PI3Ki regimen with efficacy comparable to continuous high-dose treatment but lower overall drug exposure through weekly dose modulation. This computational investigation suggests that treatment schedule optimization could reduce treatment-related toxicity while maintaining therapeutic efficacy.

In conclusion, this study establishes BRD2 as a determinant of sensitivity to PI3K/AKT inhibition and defines MSK/RSK-dependent Ser37 phosphorylation, a site absent from the other BET proteins, as a mechanism linking mitogenic and stress signaling to BRD2 chromatin occupancy. More broadly, this extends the principle that oncogenic kinase cascades act on chromatin to include acetyl-lysine readers and shows that paralog-specific regulatory sites can distinguish family members otherwise engaged indiscriminately by current inhibitors. This signaling-to-chromatin axis offers multiple points of therapeutic intervention in PI3K-driven cancers.

## Study Limitations

Several questions raised by this study remain to be addressed. Although we identify Ser37 as a functional phosphorylation site regulating BRD2 chromatin occupancy, the mechanism by which this modification alters chromatin engagement remains undefined. Phosphorylation could affect acetyl-lysine recognition, cofactor recruitment, or both. Indeed, CK2-mediated phosphorylation of BRD4 promotes its dimerization, structurally positioning the protein for chromatin recruitment and interaction with transcriptional cofactors.^79^ Determining whether BRD2 is subject to a similar mechanism will require further biochemical and structural investigation. Phosphomimetic substitution also incompletely recapitulates phosphorylation, and S37D restored BRD2 function only partially in some assays. Whether the residual phenotypes in S37D cells reflect the limitations of aspartate substitution or additional regulatory inputs at this site remains unresolved. Further, BRD4 chromatin binding is regulated by JNK kinases downstream of stress-activated MAPK signaling,^109, 110^ whereas our data implicate p38 rather than JNK in BRD2 regulation, suggesting that distinct but partially overlapping kinase inputs tune stress-responsive chromatin engagement across BET family members. Future BRD2 ChIP-seq following EGF or anisomycin stimulation will directly test how mitogenic-versus stress-induced signaling reshapes BRD2 genomic occupancy and target gene engagement. Additionally, BRD2 contains several uncharacterized ERK-targeting motifs, raising the possibility that BRD2 integrates multiple ERK-dependent inputs beyond Ser37; systematic interrogation of these sites will be needed to fully define the kinase network governing BRD2 function. Finally, current BET inhibitors are not selective among family members, so the pharmacological effects reported here cannot be attributed to BRD2 alone. BRD2-selective inhibitors or degraders will be needed to establish whether selective BRD2 targeting reproduces these effects and whether it avoids the toxicities associated with pan-BET inhibition. Finally, while the digital twin model was calibrated and validated against experimental treatment-response data, the optimized dosing regimens identified through the *in silico* clinical trials remain to be evaluated experimentally and clinically.

## Acknowledgements

We thank members of the lab for helpful discussion and critical comments on the manuscript. This work was supported by an NIH research grant CA253097 to A.T and by the Ludwig Center at Harvard to A.T, K.M.C, F.M, and T.M; an American Association for Cancer Research grant (23-40-12-HOGS), HCC SPORE in Breast Cancer Career Enhancement Program grant (NIH P50 CA168504), and NIH research grant R21CA292302 to J.H and T.M; American Cancer Society (PF-25-1379136-01-PFMBB) to TK; the Damon Runyon Cancer Research Foundation (DRQ-[2525]) to SB; and the National Science Foundation Graduate Research Fellowship under Grant No. 2140743 to J.M.

## Author Contributions

J.M. and A.T. conceived, designed, and supervised the project. J.M. performed most experiments and data analysis; A.L.H. assisted with the CRISPR/Cas9 genomic screen analysis; I.L.R. assisted with BRD2 immunoprecipitations and drug synergy analysis; A.I. and K.M.C. performed and supervised chromatin immunoprecipitation and ChIP analysis; T.K. performed cell death assays and metabolomics; J.W. and W.W. generated BRD2 knock-in cell lines; J.H. and T.M. performed and supervised breast organoid studies; J.L. and J.G.C. performed and supervised mouse studies. S.B. and F.M. developed the mathematical and pharmacokinetic models and performed the in silico clinical trial analyses. J.M., S.B., and A.T. wrote the original manuscript draft. All authors contributed to reviewing and editing of the final submission.

## STAR * Methods

### KEY RESOURCES TABLE

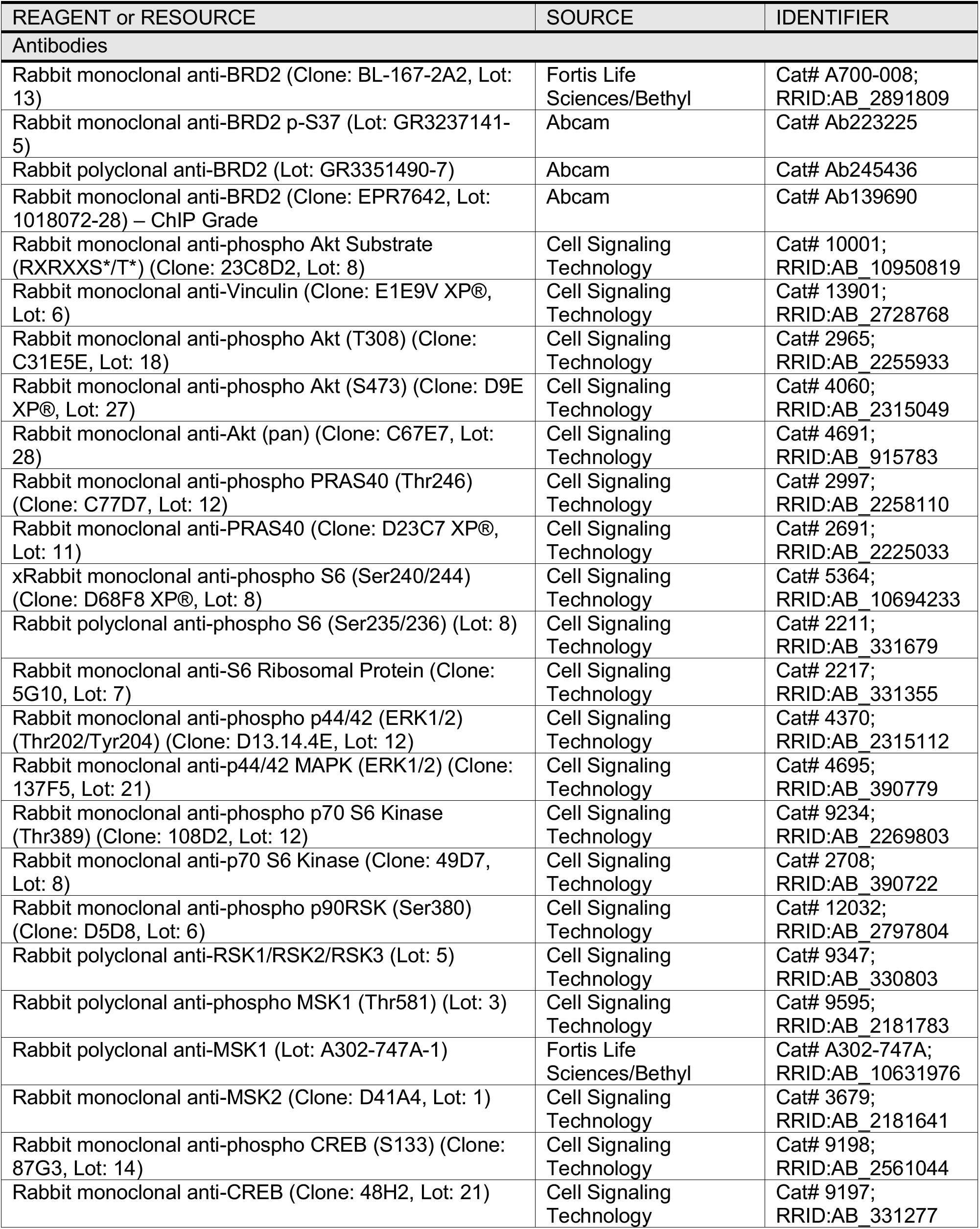
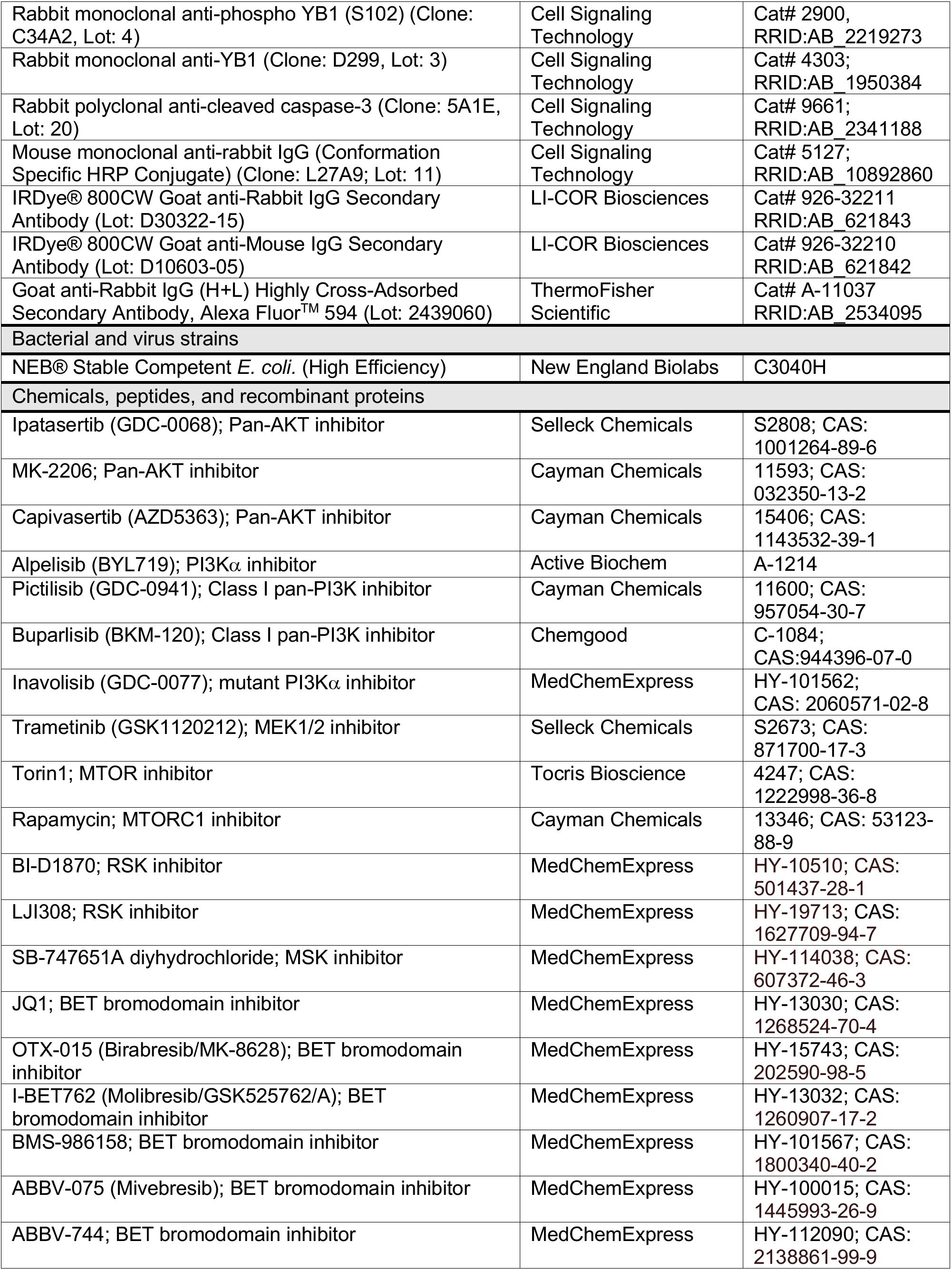
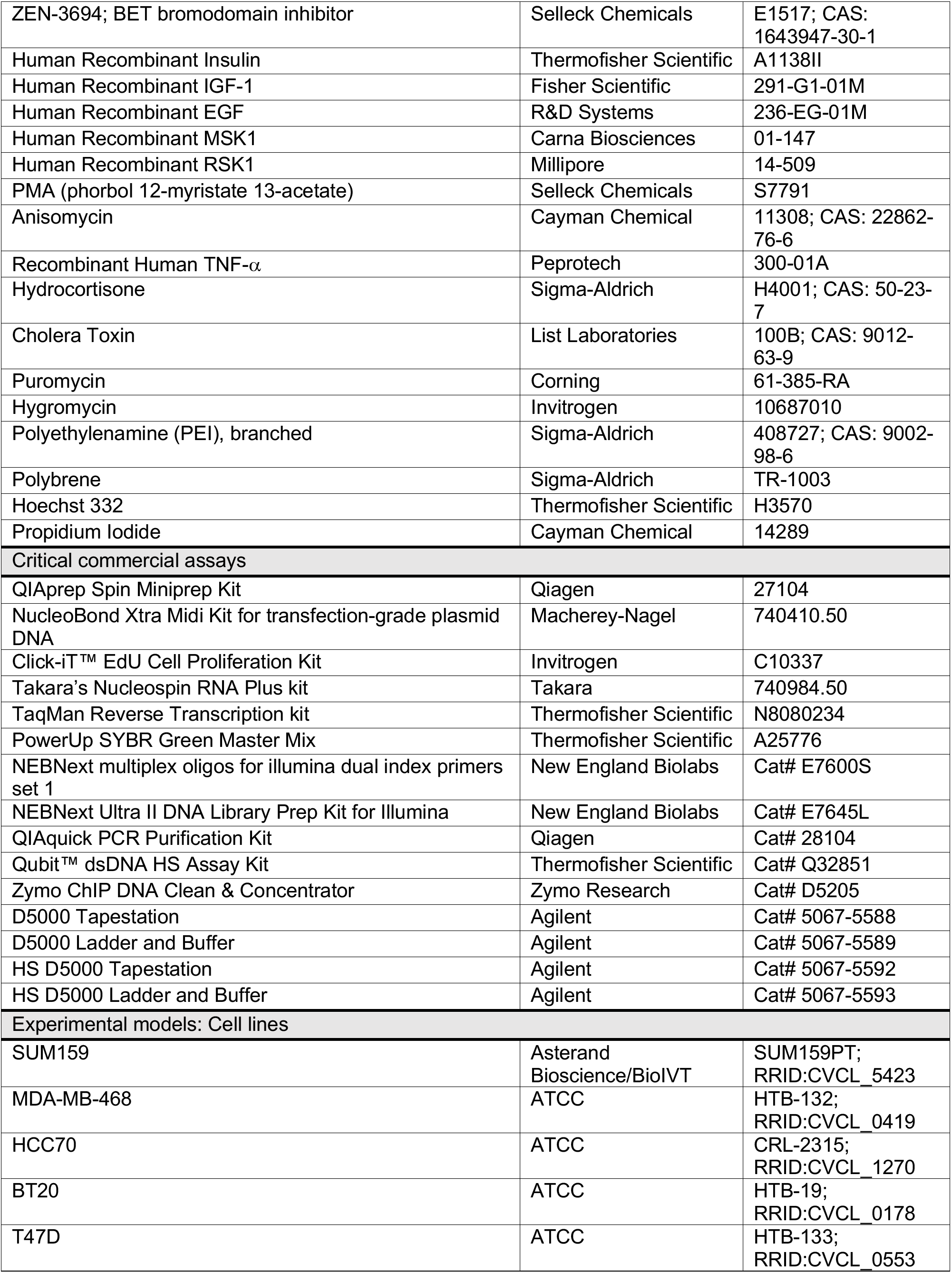
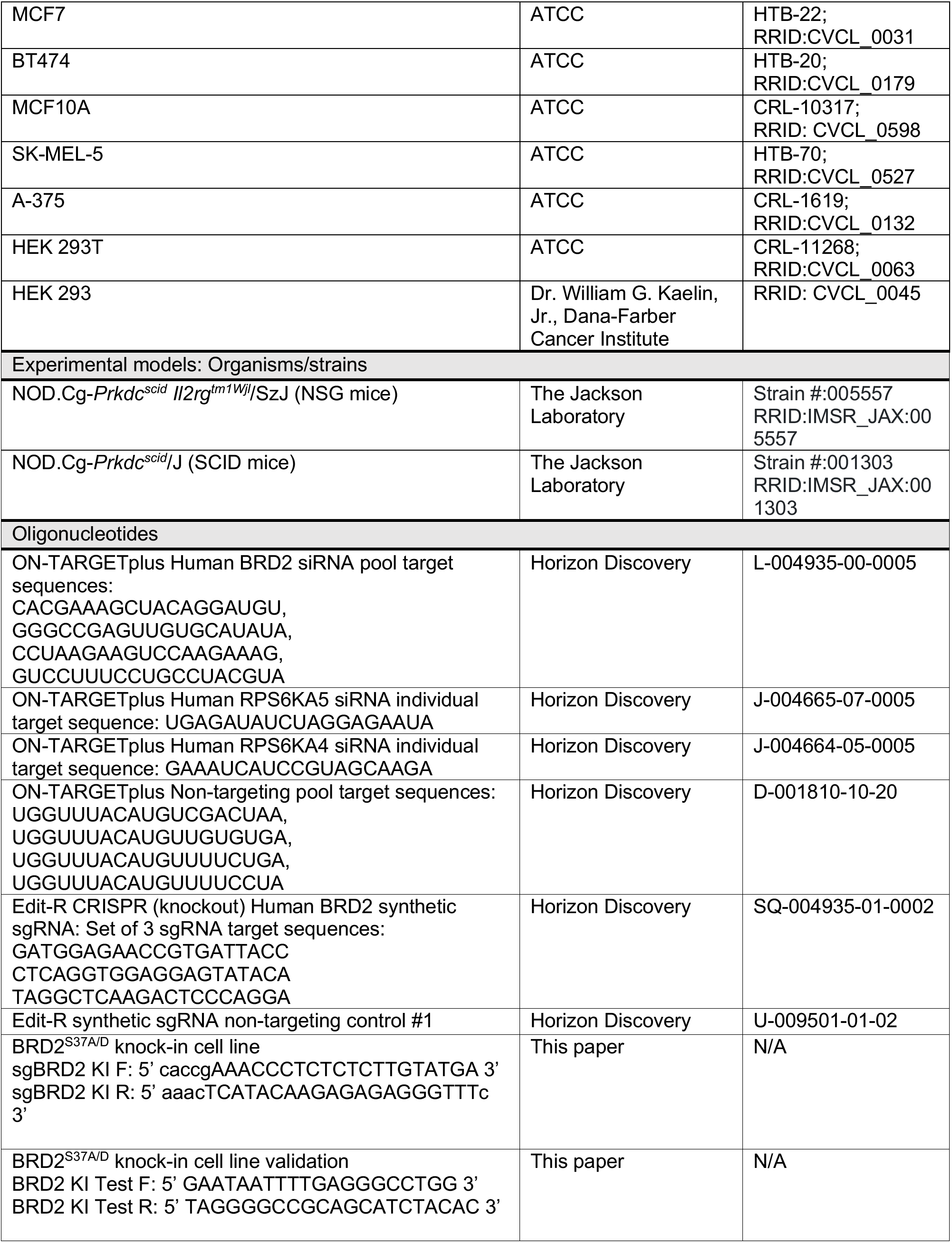
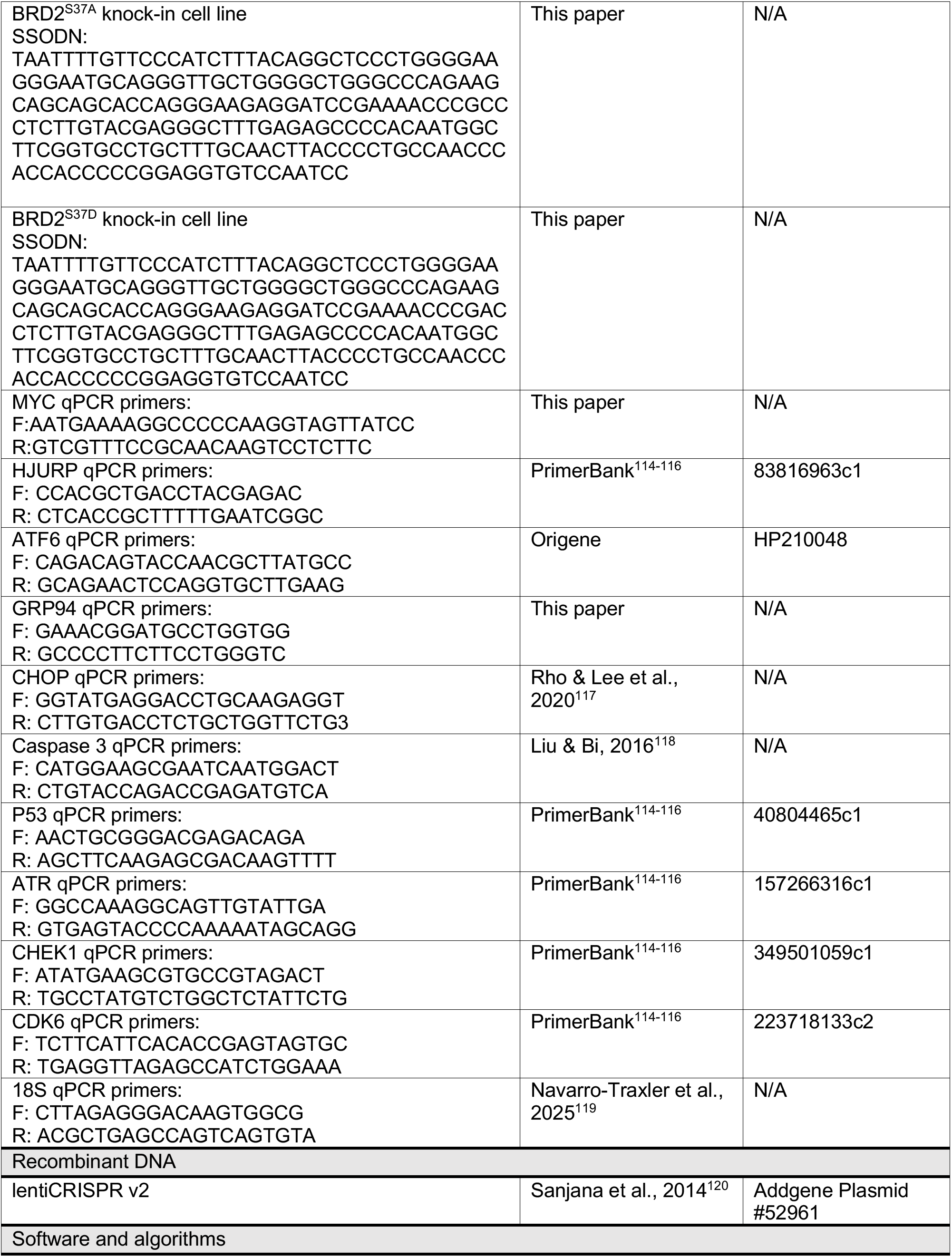
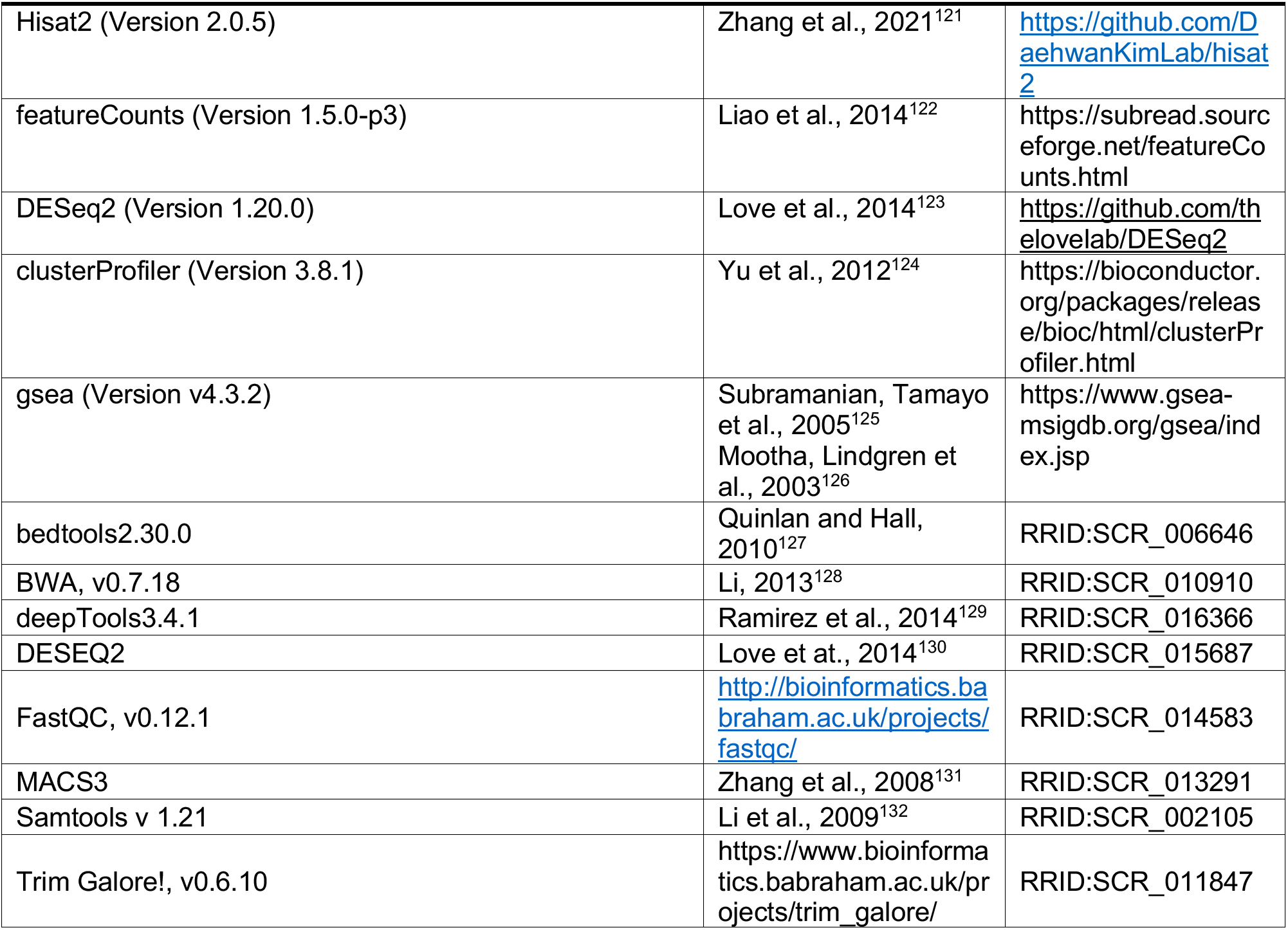

### RESOURCE AVAILABILITY

#### Lead contact

Further information and requests for resources and reagents should be directed to and will be fulfilled by the Lead Contact, Alex Toker.

#### Materials availability

All unique/stable reagents generated in this study are available from the lead contact without restriction.

#### Data and code availability

- The RNA-seq and ChIP-seq datasets generated in this study have been deposited in the NCBI Gene Expression Omnibus (GEO) database under accession codes; Drug RNA-sequencing experiment (GSE345875); KO RNA-sequencing (GSE346050), ChIP (GSE346442).
- All raw data (original western blot images and other raw data files) have been deposited to the Harvard Dataverse (https://doi.org/10.7910/DVN/MBIMTF) and are publicly available as of the date of publication. Microscopy data reported in this paper will be shared by the lead contact upon request.
- All original code has been deposited to the Harvard Dataverse (https://doi.org/10.7910/DVN/MBIMTF) and are publicly available as of the date of publication.
- Any additional information required to reanalyze the data reported in this paper is available from the lead contact upon request.

### Experimental Model and Subject Details

#### Parental Cell lines

The SUM159/SUM159PT cell line was isolated from the primary tumor of a 71-year-old female patient with ER negative, PR negative, and HER2 negative (triple-negative) anaplastic carcinoma of the breast. The MDA-MB-468 cell line was isolated from a pleural effusion of a 51-year-old Black female patient with triple-negative metastatic adenocarcinoma of the breast. The BT20 cell line was isolated from the primary tumor of a 74-year-old White female patient with triple-negative carcinoma of the breast. The HCC70 cell line was isolated from a primary ductal carcinoma of a 49-year-old Black female patient with an invasive TNM Stage IIIA grade 3 ductal carcinoma of the breast (triple-negative). The T47D cell line was isolated from a pleural effusion of a 54-year-old female patient with an infiltrating ductal carcinoma of the breast (ER positive, PR positive, and HER2 negative). The MCF7 cell line was isolated from a pleural effusion of a 69-year-old White female patient with a carcinoma of the breast (ER positive, PR positive, and HER2 negative). The BT474 cell line was isolated from a solid, invasive ductal carcinoma of the breast of a 60-year-old White female patient (ER positive, PR positive, and HER2 positive). The MCF10A cell line is an epithelial cell line isolated from the mammary gland of a 36-year-old White female patient with fibrocystic breasts (ER negative, PR negative, and HER2 negative; non-tumorigenic). The HEK293/HEK293T cell line is an epithelial cell line isolated from the embryo kidney tissue of a fetus female. The HEK293T line has been transfected to stably express the SV40 large T antigen. The SK-MEL-5 cell line was isolated from the skin tissue of a 24-year-old, White, female patient with malignant melanoma. The A375 cell line was isolated from the skin tissue of a 54-year-old, White, female patient with malignant melanoma.

#### Maintenance culture conditions

SUM159, MDA-MB-468, HCC70, T47D, MCF7, and BT474 cells were cultured in RPMI-1640 medium (Gibco, 11875093) supplemented with 10% fetal bovine serum (FBS; GeminiBio, 100-106). BT20 cells were cultured in Eagle’s Minimum Essential Medium (EMEM; Corning, 10009CV) supplemented with 10% FBS (GeminiBio, 100-106). HEK293/HEK293T, SK-MEL-5, A375 and NIH/3T3 cells were cultured in Dulbecco’s Modified Eagle’s Medium (DMEM) with L-Glutamine, 4.5 g/L glucose and sodium pyruvate (Fisher Scientific, MT10013CV) supplemented with 10% FBS (GeminiBio, 100-106). MCF10A cells were cultured in standard MCF10A growth medium without antibiotics (DMEM/F12 medium (Wisent Bioproducts, 319-075-CL), 5% horse serum (Gemini Bio, 100508), 10 µg/mL insulin (ThermoFisher Scientific/Gibco, A11382II), 500 ng/mL hydrocortisone (Sigma-Aldrich, H4001), 20 ng/mL EGF (R&D Systems, 236-EG-01M) and 100 ng/mL cholera toxin (List Biological Laboratories, 100B)). Cell lines were maintained at 37°C in a 5% CO_2_ cell culture incubator and passaged at 70-90% confluency. To passage, cells were washed once with 1X PBS and incubated for 5-10 minutes at 37°C with 0.25% Trypsin, 0.1% EDTA (Fisher Scientific, MT25053CI). Cells were passaged up to 15 times into a new dish and were maintained in culture for up to one month. Cells routinely tested negative for mycoplasma contamination every 6-9 months.

#### Modified Cell Lines

SUM159 *BRD2* KO cell lines were derived by transfection of SUM159 cells stably expressing Cas9 (transduced with lentiCRISPR v2) with pooled sgRNAs targeting *BRD2* using lipofectamine RNAiMAX transfection reagent (ThermoFisher Scientific, 13778150). Single cells were plated in 96-well plates by limiting dilution, expanded and clones were screened with via immunoblot. Clones were maintained in standard RPMI growth media with 10% FBS and 1 µg/mL puromycin.

#### Generation of the knock-in cell lines

To generate knock-in cell lines, guide RNAs (sgRNAs) targeting specific exonic regions were cloned into the lentiCRISPR v2-Puro vector (Addgene plasmid #52961). HEK293 cells were co-transfected with 500 ng of single-stranded donor DNA and 1000 ng of the sgRNA-expressing plasmid using a standard transfection protocol. Twenty-four hours post-transfection, cells were subjected to puromycin selection (1 μg/mL) for 24 hours to enrich for successfully transfected cells. Surviving cells were subsequently seeded into 96-well plates at clonal density. After approximately two weeks of expansion, individual clones were screened and validated by genomic DNA extraction followed by Sanger sequencing.

#### Transfection and lentiviral infection of plasmid DNA

HEK293T cells were transfected with lentiCRISPR v2 plasmid DNA as follows. 1.2 mL of serum-free, antibiotic-free DMEM was prepared with 11.1 µg psPAX2 (Gal/Pol), 0.6 µg VSVG, 6.3 µg of plasmid DNA.54 µL of 1 mg/mL polyethylenamine (PEI; Sigma-Aldrich, 408727) was added, and the master mix was vortexed for 10 seconds to mix and then incubated at room temperature for 15 minutes. Tissue culture-treated 10-cm dishes were coated with poly-L-lysine (PLL; Sigma-Aldrich, P1274) for 5 minutes and then washed three times with 1X PBS. HEK293T cells were trypsinized and counted, and 9 x 10^6^ cells were resuspended in 7.8 mL of DMEM + 10% FBS. The cell suspension was then added directly to the 1.2 mL transfection reaction, mixed and plated on the PLL-treated 10-cm plates. The cells were incubated for 48-72 hours before virus was harvested. Media from the HEK293T cells was passed through a 10 mL syringe with a 0.45 µm filter attached, and collected virus was aliquoted in 1 mL stocks and stored at −80°C. Target cell lines were seeded in 6 cm plates at 75,000-100,000 cells/mL in complete media (RPMI-1640 + 10% FBS). 24 hours after seeding, media was aspirated from the target cell line and 1 mL of virus was added to each plate. Polybrene (Sigma-Aldrich, TR-1003) was added at a final concentration of 5 µg/mL, and cells were incubated at 37°C for 4 hours. After which 3 mL of complete media was added to each plate, and cells were incubated at 37°C for 48 hours. Media was then changed on the transduced cells to antibiotic containing media (2 µg/mL puromycin). Cells were maintained and expanded in antibiotic-containing media.

#### CRISPR/Cas9 genome editing

SUM159 cells stably expressing Cas9 were first generated by lentiviral transduction of the lentiCRISPR v2 plasmid (See Transfection and lentiviral infection of plasmid DNA details above). These cells were then transfected with 25 nM of Edit-R synthetic sgRNAs (either non-targeting, or a pool of 3 *BRD2* targeting sgRNAs) using lipofectamine RNAiMAX transfection reagent (ThermoFisher Scientific, 13778150) as follows. Stocks of guides were prepared by resuspending sgRNAs using nuclease-free 10 mM Tris-HCl Buffer pH 7.4 (PerkinElmer/Horizon Discovery, B-006000-100) for a stock concentration of 10 µM. Following addition of buffer, the solution was placed on an orbital mixer for 30 minutes at room temperature and then aliquoted into microcentrifuge tubes and stored at −20C. Each aliquot was freeze-thawed no more than five times. For sgRNA transfection in 6 well dishes, cells were seeded at a density that gives 70-90% confluency the next day (for Cas9 expressing SUM159 cells: 25,000 cells/ml). 24 hrs after seeding all sgRNAs were diluted to a working concentration of 2 µM in 10 mM Tris-HCl Buffer pH 7.4 and further diluted in 250 µL of pre-warmed OptiMEM (ThermoFisher Scientific/Gibco, 31985062) in tube A so that the final concentration on cells would be 25 nM total. Tube B was prepared by diluting Lipofectamine RNAiMAX transfection reagent (ThermoFisher Scientific/Invitrogen, 13778075) in 250 µL of pre-warmed OptiMEM so that the final concentration would be 2.5 µL of Lipofectamine RNAiMAX for every 1 mL of final media. Tubes A and B were each mixed individually by pipetting and incubated at room temperature for 5 minutes. Tube A was thoroughly mixed with Tube B, incubated at room temperature for 20 min, and 400 µL of the mixture was added dropwise to cells containing 1.6 mL of antibiotic-free media. Cells were incubated with the sgRNA/transfection reagent for 72 hrs before expansion to a 10 cm dish. Once plate was confluent, cells were plated for initial western validation of knockout in pooled population, and as single cells in 96 well plates (seeded cells at 5 cells/mL in 100 uL/well). Single cell clones were grown in standard RPMI growth media with 10% FBS and 1 ug/mL puromycin and were screened for BRD2 expression via immunoblot.

#### Growth factor/Inhibitor treatments for immunoprecipitation (IP) of BRD2

MCF10A cells were serum/growth factor deprived for 18 hours before pre-treatment with inhibitors indicated (Inhibitor information listed in Key Resources Table) for 30 minutes. Inhibitor stocks were dissolved in DMSO at 10 mM under sterile conditions, aliquoted, and stored at −80°C. Aliquots were freeze-thawed no more than five times. After pre-treatment, cells were stimulated with a growth factor (either 100 nM Insulin (ThermoFisher Scientific / Gibco, A11382II; stock: 100 µM in 0.5M HCL, then diluted in water), 50 ng/mL IGF-1 (Fisher Scientific, 291-G1-01M; 100 µg/ml in 0.1% (w/v) BSA/PBS), 10 ng/mL EGF (R&D Systems, 236-EG-01M; stock 100 µg/ml in 0.1% (w/v) BSA/PBS), 100 nM PMA (phorbol 12-myristate 13-acetate) (Selleckchem, S7791; stock: 100 µM in DMSO), 10 µM Anisomycin (Cayman Chemical, 11308; stock 20 mM in DMSO), or 10 ng/mL TNF-α (Peprotech, 300-01A; stock 10 µg/ml in 0.1% (w/v) BSA/PBS)) for 30 minutes or otherwise indicated time period. All growth factor stocks were prepared under sterile conditions, aliquoted, and stored at −80°C.

#### Immunoprecipitation (IP) of BRD2

All steps were carried out on wet ice or at 4°C. Gentle protein harvest protocol was followed as previously described.^133^ Cells in 10 cm dishes were lysed in 600 µl of NP40 lysis buffer [1X stock stored at 4°C: 40 mM HEPES, pH 7.5 (ThermoFisher Scientific / Gibco, 15630080); 120 mM NaCl (Sigma-Aldrich, S7653); 1 mM EDTA, pH 8.0 (Sigma-Aldrich, E6758); 1% IGEPAL CA-630 (NP40) (Sigma-Aldrich, I3021); 10 mM sodium pyrophosphate tetrabasic (Sigma-Aldrich, S6422); 10 mM β-glycerophosphate (Sigma-Aldrich, G9422S); 50 mM sodium fluoride (Fisher Scientific, S25547); 5% (v/v) glycerol (Fisher Scientific, G33-500)] with 1:100 protease inhibitor cocktail [104 mM AEBSF, 80 µM Aprotinin, 4 mM Bestatin, 1.4 mM E-64, 2 mM Leupeptin, and 1.5 mM Pepstatin A (Sigma-Aldrich, P83400)] added day of use. Plates were scraped into 1.5 ml tubes and incubated on a nutator (Clay Adamas, 421105) at 4°C for 20 minutes before centrifugation at 15,000 x g for 10 minutes at 4°C. Cleared supernatants were transferred to a new tube and were frozen at −20°C and thawed once to lyse cellular nuclei before running protein quantification. Protein content was measured by DC protein assay (based on Lowry assay; Bio-Rad, 5000116). Lysates were normalized for protein concentration and samples were brought up to the same volume (at least 500 µl) in NP40 lysis buffer so that at least 500 µg of protein was used for the IP of BRD2. 50 µl of lysate was saved for immunoblotting (lysate was normalized for immunoblotting for whole cell lysate: WCL as described below). Lysate normalized for IP was incubated for 2 hours with 2.5 µl of BRD2 IP antibody (Fortis Life Sciences/Bethyl, A700-008), rocking on a nutator (Clay Adamas, 421105) at 4°C. Protein G Dynabeads (ThermoFisher Scientific, 10004D) were aliquoted into a 1.5 ml Eppendorf tube and washed once with bead wash buffer (40 mM HEPES, pH 7.5; 120 mM NaCl). Initial bead wash completed by placing an Eppendorf tube containing aliquoted Dynabeads on DynaMagTM-2 magnet (ThermoFisher Scientific, 12321D) for one minute and discarding the supernatant by aspiration with a pipette, before resuspending the beads in 1 mL of bead wash buffer by inverting 8-10 times. The tube was then again placed on the magnet, bead wash buffer was discarded, and beads were resuspended in a 1:1 slurry of NP40 lysis buffer. 20 µl of resuspended beads was added to each IP sample for an additional 2 hours, rocking at 4°C. IP samples were then placed on magnet for 1 minute and the supernatant was discarded before sample beads were washed three times in NP40 lysis buffer (wash in 500 µl NP40 lysis buffer, invert 8-10 times to resuspend beads, incubate on nutator (Clay Adamas, 421105) at 4°C for 10 minutes). Following washes, sample beads were allowed to precipitate on magnet, the supernatant was discarded, and sample beads were resuspended in 40 µl of 1X Laemmli sample buffer [1X: 50 mM Tris Base, pH 6.8 (Fisher Scientific, BP152); 2% (w/v) Sodium Dodecyl Sulfate (SDS) (Fisher Scientific, NC0755841); 5% (v/v) β-mercaptoethanol (Sigma-Aldrich, M3148); 5% (v/v) glycerol (Fisher Scientific, G33-500); 0.01% (w/v) Bromophenol Blue; (6X stock aliquots stored at −20°C and freeze-thawed no more than three times)]. Samples were heated at 95°C for 5 minutes and stored at −20°C.

#### Immunoblotting (for BRD2 IP)

BRD2 IP samples were heated at heated at 95°C for 5 minutes and run by SDS-PAGE on 10% acrylamide gels in 1X running buffer (Boston BioProducts, BP-150), or 4-12% Bis-Tris gradient gels (ThermoFisher Scientific, NP0329BOX) in MOPS running buffer (ThermoFisher Scientific, NP000102). Proteins were transferred to PVDF membranes (Millipore-Sigma, IPVH00010) at a constant 300 mA for 90 minutes at 4°C in Tris-glycine transfer buffer [10X Transfer (Electro) Blotting Buffer (Boston BioProducts, BP-190) diluted to 1X in water with 10% (v/v) methanol]. Membranes were blocked with 5% (w/v) bovine serum albumin (BSA; Gold Biotechnology, A-420-100) in tris-buffered saline (TBS; Boston BioProducts, BM-301), rocking for 1 hour at room temperature. Membranes were incubated in primary antibody diluted in 5% (w/v) BSA in TBST with 0.01% 0.01% (w/v) sodium azide (Fisher Scientific, S227I-25), rocking at 4°C overnight. Membranes were washed 3 x 10 min in TBST at room temperature and incubated in horse radish peroxidase (HRP)-conjugated secondary antibody diluted in 5% (w/v) non-fat dry milk (Fisher Scientific, NC9022655) in TBST for 1 hour, rocking at room temperature. Membranes were washed 3 x 10 min TBST at room temperature and incubated in Clarity Western ECL Substrate (Bio-Rad, 1705061) for 1-2 min. The chemiluminescent signal was detected using film (Thomas Scientific, F-BX810).

#### Immunoblotting (WCL)

Cells were washed once in cold 1X PBS (Boston BioProducts, BM-220) and collected on wet ice in 4°C RIPA lysis buffer (150 mM Tris-HCl, 150 mM NaCl, 0.5% (w/v) sodium deoxycholate (Sigma-Aldrich, D6750), 1% (v/v) NP-40 (Sigma-Aldrich, I3021), pH 7.5) containing 0.1% sodium dodecyl sulfate (SDS; AmericanBio, AB01920-00500), 1 mM sodium pyrophosphate (Na_4_P_2_O_7_; Sigma-Aldrich, S390-500), 20 mM sodium fluoride (NaF; Fisher Scientific, S25547), 0.5% (v/v) protease inhibitor cocktail (104 mM AEBSF, 80 μM aprotinin, 4 mM bestatin, 1.4 mM E-64, 2 mM leupeptin and 1.5 mM pepstatin A; Sigma-Aldrich, P8340) and 50 nM calyculin A (LC Laboratories, C-3987), added just before use. Plates were scraped into 1.5 mL microcentrifuge tubes, vortexed, and incubated on wet ice for 15 minutes. Samples were then centrifuged at 14,000 rpm for 10 minutes at 4°C. Cleared supernatants were transferred to a new tube and protein content was measured by DC protein assay (based on Lowry assay; Bio-Rad, 5000116). Sample concentrations were normalized using 2X SDS sample buffer (62.5 mM Tris pH 6.8, 2% SDS, 10% glycerol (Fisher Scientific G33-500), bromophenol blue (Fisher Scientific, BP115-25), 5% (v/v) β-mercaptoethanol (Sigma-Aldrich, M3148)). Cell lysates were boiled at 95°C for 5 minutes and stored at −20°C. Cell lysates were run by SDS-PAGE on 7.6-15% acrylamide gels in 1X running buffer (Boston BioProducts, BP-150). Proteins were transferred to nitrocellulose membranes at 100 volts for 90 minutes in Tris-glycine transfer buffer [10X Transfer (Electro) Blotting Buffer (Boston BioProducts, BP-190) diluted to 1X in water with 10% (v/v) methanol]. Membranes were blocked with 5% (w/v) bovine serum albumin (BSA; Gold Biotechnology, A-420-100) in tris-buffered saline (TBS; Boston BioProducts, BM-301), rocking for at least 1 hour at room temperature. Membranes were incubated in primary antibody diluted in 5% (w/v) BSA in TBST with 0.01% 0.01% (w/v) sodium azide (Fisher Scientific, S227I-25), rocking at 4°C overnight. Membranes were washed 3 x 10 min in TBST at room temperature and incubated for 1 hour rocking at room temperature with fluorophore-conjugated secondary antibodies (LI-COR Biosciences). Membranes were washed two times for 10 minutes each in TBST and one time for 10 minutes in TBS and imaged with the Odyssey M Imaging System (LI-COR Biosciences).

#### In vitro kinase assay

For the *in vitro* kinase assay, BRD2 immunoprecipitations using lysates from 10 cm dishes (of HEK293 WT or S37A knock-in cells) were performed as described above until the final resuspension step. After three washes in NP40 lysis buffer, two additional washes were performed in 500 µL kinase reaction buffer [20 mM, HEPES, pH 7.5 (ThermoFisher Scientific / Gibco, 15630080); 10 mM MgCl_2_ (Sigma, M2670-100G); 0.5 mM EGTA (Sigma, E4378-25G)]. All supernatant was removed, and the beads were resuspended in 20 µL kinase reaction buffer with 100 µM ATP (Sigma, A2383-1G) and 1 mM DTT (Fisher, BP172-5) with or without 50 µM recombinant MSK1 or RSK1 kinases [provided as a gift from Johnson et al.^82^; MSK1 (Carna Biosciences, 01-147: LOT 08CBS-0491H); RSK1 (Millipore, 14-509: LOT 194681-B)]. Samples were incubated for 30 minutes at 32°C in a thermomixer (Eppendorf, 5382000023) with 250 rpm agitation. Reaction was stopped on ice, and a portion of each sample supernatant was saved with sample buffer added to 1X final concentration. The beads were resuspended in 1X Laemmli sample buffer. Samples were heated at 95°C for 5 minutes and stored at −80 °C. For immunoblotting, ¼ of each immunoprecipitation sample was loaded per SDS-PAGE gel.

#### RNA interference using siRNAs

Stocks of small interfering RNAs (siRNAs) were prepared by resuspending siRNAs using 1X siRNA buffer [5X siRNA buffer (PerkinElmer/Horizon Discovery, B-002000-UB-100), diluted to 1X in sterile molecular grade RNase-free water (PerkinElmer/Horizon Discovery, B-12-003000-WB-100)] for a stock concentration of 20 µM. Following addition of buffer, the solution was placed on an orbital mixer for 30 minutes at room temperature and then aliquoted into microcentrifuge tubes and stored at −20C. Each aliquot was freeze-thawed no more than five times. For siRNA treatments in 10 cm dishes, siRNAs were diluted in 500 µL of pre-warmed OptiMEM (ThermoFisher Scientific/Gibco, 31985062) in tube A so that the final concentration on cells would be 25 nM total. Tube B was prepared by diluting Lipofectamine RNAiMAX transfection reagent (ThermoFisher Scientific/Invitrogen, 13778075) in 500 µL of pre-warmed OptiMEM so that the final concentration would be 1.5 µL of Lipofectamine RNAiMAX for every 25 µL of Opti-MEM media. Tubes A and B were each mixed individually by pipetting and incubated at room temperature for 5 minutes. Tube A was thoroughly mixed with Tube B, incubated at room temperature for 20 min, and added dropwise to cells. Cells were incubated with the siRNA/transfection reagent overnight and media was replaced with fresh growth media the next day. Approximately 30 hours post-transfection, cells were washed with serum/growth-factor-free media, and serum/growth factor starved for 18 hours. The cells were then used in the growth factor/inhibitor experiments described above.

#### Cell density (Sulforhodamine B) assays

Cells were seeded in tissue culture-treated 96-well plates in 90 µL of appropriate growth media (SUM159: 2000 cells/well, MDA-MB-468: 5000 cells/well, BT20: 6000 cells/well, HCC70: 6000 cells/well, T47D: 6000 cells/well, MCF7: 4000 cells/well, BT474: 6000 cells/well, MCF10A: 2000 cells/well, SK-MEL-5: 4000 cells/well, A375: 4000 cells/well, HEK293: 3000 cells/well). After 24 hours, Day 0 cells were fixed by addition of trichloroacetic acid directly to the media to a final concentration of 8.33% (w/v). Experimental cells were then treated with 5-10 µL of drug to bring the final volume in each well to 100 µL. Cells were then fixed at determined endpoints of treatment for each experiment (Day 1, 2, 3, and/or 4). Cell density was assayed using sulforhodamine B (SRB; Sigma-Aldrich, 230162) staining, as previously described.^134^ Relative cell density was determined at each time point by normalizing to the day 0 control. For dose curve and double dose curve experiments, these values were normalized from 0-100 using GraphPad PRISM, where an empty well (background) served as the 0% reference, and untreated cells served as the 100% reference. Normalized cell densities were plotted versus log10 drug concentration, and a nonlinear curve was fit using the log(inhibitor) vs. normalized response -- Variable slope function in GraphPad PRISM. IC_50_ values were calculated by GraphPad PRISM based on the nonlinear curve fit. Cell density data for experiments that were not dose curves or double dose curves were transformed to log2(Y) values using GraphPad PRISM and plotted such that values less than 0 indicate cytotoxicity or cell death.

#### Synergy calculations

For proliferation assays with two inhibitors, synergy scores were calculated using the synergyfinder R package.^135^ HSA synergy scores were reported for each drug dose combination tested and displayed as a heatmap.

#### PI death assay

HCC70 and T47D were seeded in 96-well plates and allowed to adhere overnight. Cells were then treated with GDC-0077 (20 µM for HCC70; 500 nM for T47D), ZEN-3694 (40 µM), or their combination for 72 hrs. Cells were then incubated with 3 µg/ml Hoechst 3342 (ThermoFisher, H3570), and 1 µg/ml propidium iodide (Cayman Chemical, 14289) at 37 °C and imaged using a Celigo Imaging Cytometer. Two images were taken per well and percent cell death was calculated as the ratio of PI-positive cells to total Hoechst-positive cells.

#### Real-time quantitative polymerase chain reaction (RT-qPCR)

Total RNA was extracted from cells using the NucleoSpin RNA Plus kit (Macherey-Nagel, 740984) following the manufacturer’s instructions. Reverse transcription was performed with the TaqMan Reverse Transcription kit (ThermoFisher Scientific, N8080234) to generate cDNA. For quantitative PCR, cDNA was amplified using PowerUp SYBR Green Master Mix (TheromoFisher Scientific, A25776). A reaction master mix was prepared containing 250 nM forward primer, 250 nM reverse primer, 1 X SYBR Green Master Mix, and nuclease-free water to a final volume of 10 µL per well. Experiment was performed in a 384-well plate, with each reacting having 10 µL of master mix and 2 µ of cDNA at 2.5 ng/µL (5 ng total cDNA per reaction). Plates were briefly centrifuged at 1000 rpm to ensure entire volume was at the bottom of the well. RT-qPCR was performed on a CFX384 Touch Real-Time PCR Detection System (Bio-Rad) with the following cycling conditions: 50 °C for 2 min, 95 °C for 2 min, followed by 40 cycles of 95 °C for 15 s and 60 °C for 1 minute, with a final melt curve at 65 °C for 5 s and 95 °C for 5 s. All reactions were run in technical triplicate. Relative mRNA expression was calculated using the ΔΔCT method, normalizing to 18s ribosomal RNA. Primers (listed in **Key Resources Table**) were either designed using NCBI Primer-BLAST or from the PrimerBank and validated to have efficiencies between 90-110% in each cell line.

#### Mouse Studies

All animal experiments were performed at Beth Israel Deaconess Medical Center (BIDMC) in accordance with the guidelines of the BIDMC Institutional Animal Care and Use Committee (IACUC). *BRD2 KO Xenograft Experiment:* Non-targeting or *BRD2* KO generated SUM159 cells (described above) were maintained in RPMI-1640 supplemented with 10% FBS + 1 µg/ml Puromycin, and cells tested negative for mycoplasma before injection. On the day of injection, the cells were washed twice with 1X PBS, trypsinized and counted. A total of 5 x 10^6^ cells per mouse were resuspended in 100 µL of serum-free RPMI-1640 and placed on wet ice. Cells were mixed with Matrigel (Corning, 356230) in a 1:1 (v/v) ratio and injected orthotopically into the mammary fat pad of 20 NSG mice (10 injected with NT and 10 with *BRD2* KO). Tumors were measured with calipers 2 times per week (length and width) until tumors reached the humane endpoint. After 47 days of treatment, all mice injected with SUM159 non-targeting were euthanized. Sections of tumor, liver, and kidney were snap frozen and fixed in 10% formalin for immunoblotting and immunohistochemistry, respectively. *Drug Xenograft Experiments:* SUM159 or HCC70 cells tested negative for mycoplasma before injection. On the day of injection, the cells were washed twice with 1X PBS, trypsinized and counted. A total of 5 x 10^6^ cells per mouse were resuspended in 100 µL of serum-free RPMI-1640 and placed on wet ice. Cells were mixed with Matrigel (Corning, 356230) in a 1:1 (v/v) ratio and injected orthotopically into the mammary fat pad of NSG mice.

Tumors were allowed to grow for 16-18 days before mice were assigned to each treatment group (vehicle, GDC-0077, ZEN-3694, combination) and treatments were administered daily by oral gavage. GDC-0077 (Genetech) was prepared as a suspension in 0.5% carboxymethylcellulose/0.2% Tween-80 and was dosed daily at 5, 10, 15, or 25 mg/kg depending on the experiment. ZEN-3694 (Zenith Epigenetics) and was dosed daily at 25, 50, 75, or 100 mg/kg depending on the experiment. ZEN-3694 was prepared in 0.5% carboxymethylcellulose/0.2% Tween-80, 95% corn oil/5% DMSO or formulation EA006. Vehicle EA006 (Polyethylene Glycol-300, 10% v/v; Polysorbate-80, 2.5% v/v; sterile water to final volume) was prepared fresh at the start of each experiment and filter sterilized. ZEN-3694 was prepared fresh daily in the recommended EA006 formulation. Briefly, weighed ZEN-3694 powder was dissolved in 5 times the volume of 1N HCL to make an in-situ HCL salt. Dissolved ZEN-3694 slurry was then mixed with Polyethylene Glycol-300, 10% v/v; Polysorbate-80, 2.5% v/v, and sterile water to final volume. Both ZEN-3694 and GDC-0077 was sonicated twice for 5-minute bursts, and vortexed vigorously before administration. Tumors were measured with calipers 2 times per week (length and width), and all mice in each experiment were euthanized when tumors in any condition reached humane endpoint. Mice in each group were treated with GDC-0077 (2 hours) or ZEN-3694 (6 hours) prior to euthanasia and sections of tumor, liver, and kidney were snap frozen and fixed in 10% formalin for immunoblotting and immunohistochemistry, respectively.

#### Patient-derived organoid cultures

Propagation and culturing of patient-derived organoid cultures (PDOs) was previously described.^136^ Briefly, PDOs were incubated in 1× Dispase-II solution with 2 mg/mL collagenase for 30-45 minutes at 37°C and mechanically disrupted by passing through a 26G needle.PDOs were washed once with Advanced DMEM/F12 supplemented with 5% FBS and pelleted by centrifugation at 400 x g for 5 minutes. PDO fragments were embedded in Cultrex growth factor-reduced basement membrane extract type II (Fisher Scientific, 35-330-1002), 50 µL drops were plated into a 24-well plate, and 500 µL PDO media was added 30 minutes later.^136^

To assess drug sensitivity, 200-600 PDO fragments were plated into 8-well chamber slides, and 1 μM AZD5363 (Cayman Chemicals, 15406), 1 μM GDC-0077 (MedChemExpress, HY-101562) and/or 3 µM ZEN-3694 (Selleck Chemicals, E1517) was added the following day. After 96 hours of drug treatment, PDOs were pulsed with 10 μM 5-ethynyl-2′-deoxyuridine (EdU) for 4 hours and fixed with 4% paraformaldehyde for 30 minutes. To assess cell proliferation and apoptosis, fixed PDOs were permeabilized with wash buffer (0.3% Triton X-100 in PBS) for 20 minutes. EdU labeling was performed for 40 minutes using the EdU Click-IT imaging kit (Invitrogen, C10337) according to manufacturer’s description. PDOs were washed 3 times with wash buffer, blocked for 1 hour with blocking buffer (5% goat serum, 0.2% BSA, 0.3% Triton X-100 in PBS) and incubated with anti-cleaved caspase-3 (Cell Signaling Technology, 9661) in blocking buffer overnight at 4°C. The following day the PDOs were washed extensively with wash buffer, incubated with secondary antibody (Alexa Fluor 488) for 2 hours at room temperature, washed with wash buffer and mounted using Vectashield mounting media containing DAPI (Vector Laboratories, H-1200-10). PDOs were imaged with Zeiss LSM 880 confocal microscope. To assess proliferation, 10-20 PDOs per treatment condition were imaged and the ratio of EdU positive cells per total number of cells was quantified using original images. To assess apoptosis, 10-20 PDOs per treatment condition were scored based on the presence of cleaved caspase-3 staining using original images. Plotted representative images were normalized to have equal brightness using ImageJ/Fiji.

#### Patient-derived organoid characteristics

| Patient | Pathology | Mutations |
| --- | --- | --- |
| 8 | ER-positive metastatic breast cancer (bone) | *PIK3CA* H1047R |
| 10 | ER-low metastatic breast cancer (ascites) | *AKT* E17K |
| 26 | ER-positive metastatic breast cancer (bone) | - |

#### RNA-sequencing

##### For BRD2 KO Experiment

SUM159 non-targeting and *BRD2* KO cells (described above) were plated at 100,000-250,000 cells/ml in RPMI-1650 supplemented with 10% FBS + 1 µg/ml Puromycin in 6cm tissue cultured plates to achieve 75% density. The next day, each plate was washed once with 1 ml of cold 1X PBS and aspirated completely. Plates were snap frozen on dry ice and stored at −80°C until all biological replicates were collected. Four biological replicates were seeded on sequential days and harvested at the same time of day. Samples were collected for protein harvest in parallel to confirm expected *BRD2* KO. Snap frozen plates were thawed on ice, and RNA was extracted using Takara’s Nucleospin RNA Plus kit (Takara, 740984.50). RNA quantity and purity were assessed by Nanodrop 1000. Samples were submitted to Novogene for integrity assessment (Agilent 2100 analysis), mRNA library preparation (unstranded) and paired-end (150 bp) sequencing on a NovaSeq6000, S4 flow cell. *For Combination Drug Experiment:* SUM159 cells were plated at 100,000-250,000 cells/ml in RPMI-1640 supplemented with 10% FBS in 6cm tissue cultured plates to achieve 75% density at the end of drug treatment. The next day, cells were treated with 1 µM GDC-0077 for 24hrs. Six hours before the 24-hour mark, cells were treated with 5 µM ZEN-3694. After 24hrs of either DMSO vehicle, single agent PI3Ki, six hours of single agent BETi, or their combination, each plate was washed once with 1 ml of cold 1X PBS and aspirated completely. Plates were snap frozen on dry ice and stored at −80°C until all biological replicates were collected. Four biological replicates were seeded on sequential days and harvested at the same time of day. Samples were collected for protein harvest in parallel to confirm the inhibitors on-target efficacy. Snap frozen plates were thawed on ice, and RNA was extracted using Takara’s Nucleospin RNA Plus kit (Takara, 740984.50). RNA quantity and purity were assessed by Nanodrop 1000. Samples were submitted to Novogene for integrity assessment (Agilent 2100 analysis), mRNA library preparation (unstranded) and paired-end (150 bp) sequencing on a NovaSeq6000, S4 flow cell.

#### RNA-Sequencing Analysis

RNA-sequencing data analysis in this study was completed utilizing Novogene’s bioinformatic analysis pipeline, which is described briefly below. *Data quality control:* Raw data (raw reads) in fastq format was first processed through fastp software. In this step, clean data (clean reads) were obtained by removing reads containing adapter, ploy-N and low-quality reads. Q20, Q30 and GC content of the clean data were calculated. All downstream analyses were based on the clean data with high quality. *Read mapping to the reference genome and quantification of gene expression level:* Reference genome and gene model annotation files were downloaded from genome website. Index of the reference genome was built using Hisat2 v2.0.5 and paired-end clean 1 reads were aligned to the reference genome using Hisat2 v2.0.5. featureCounts v1.5.0-p3 was used to count the reads numbers mapped to each gene. The FPKM (Fragments Per Kilobase of transcript sequence per Millions base pair sequenced) of each gene was calculated based on the length of the gene and read counts mapped.

##### Differential Expression Analysis

Differential expression analysis for two groups was performed using the DESeq2 R package (1.20.0). This package provided statistical programs for determining differential expression in digital gene expression data using models based on negative binomial distribution. The resulting P-values were adjusted using the Benjamini and Hochberg’s methods to control the error discovery rate. The corrected P-value ≤ 0.05 & |log2(foldchange)| ≥ 1 was set as the threshold of significant differential expression. *Enrichment Analysis:* Gene Ontology (GO) enrichment analysis of differentially expressed genes was implemented by the clusterProfiler R package (3.20) in which gene length bias was corrected. In addition, this package was utilized to test the statistical enrichment of differential expression genes in KEGG, Reactome, DO (Disease Ontology), DisGeNET databases. All terms with corrected Pvalue less than 0.05 were considered significantly enriched by differential expressed genes. Gene Set Enrichment Analysis (GSEA) waws performed using a local version of the GSEA analysis tool http://www.broadinstitute.org/gsea/index.jsp. GO, KEGG, Reactome, DO, and DisGeNET data sets were used for GSEA independently. ssGSEA was performed using GenePattern (https://www.genepattern.org/).^125, 137^ Gene sets are published Hallmark, KEGG, and Reactome gene signatures.^68, 111–113, 138^

#### Chromatin Immunoprecipitation Sequencing (ChIP-Seq)

HEK293 BRD2 knockin cells (WT, KO, S37A, and S37D; 10 x 10^6^ per ChIP) were cross-linked with 1% formaldehyde (Sigma, #252549) for 5 minutes at room temperature with mild rotation, and the reaction was quenched with 0.125 M glycine for 5 minutes. Cells were washed twice with cold PBS, pelleted, and snap-frozen at −80°C. Cell pellets were resuspended in ChIP buffer (150mM NaCl, 1% TritonX-100, 5mM EDTA, 10mM TrisCl, pH 7.5, 0.5 mM DTT) containing protease inhibitors (Pierce, #A32963), 0.5 mM DTT, and SDS was added to a final concentration of 0.5%. Chromatin was sheared using a Covaris E220 sonicator (4 mL milliTUBEs, PN 520135; peak power 140.0, duty factor 5.0%, 200 cycles/burst, average power 7.5 W, 4°C) for approximately 20 min per sample. Fragment sizes were verified to fall predominantly between 200-900 bp by Qubit and Tapestation (Agilent D500/HSD5000) analysis following proteinase K/RNAse digestion and reversal of cross-links. Sheared chromatin was cleared by centrifugation (13,000 rpm, 10 min, 4°C) and diluted in ChIP buffer (with protease inhibitors and DTT, no SDS) to reduce SDS to 0.1-0.15%. An input aliquot was reserved. Immunoprecipitation was performed by incubating diluted chromatin with the appropriate antibody (BRD2; Abcam: ab139690; H3K27ac; Abcam: ab4729; IgG; Cell Signaling Technology: #2729) and 120 µL Protein A/G Dynabeads (Thermofisher, #10004D) overnight at 4°C with rotation. Beads were washed sequentially with Mixed Michelle Wash Buffer (150mM NaCl, 20mM TrisCl, pH 8.0, 5mM EDTA, 5.2%. sucrose, 1% TritonX-100, 0.2% SDS), Buffer 500 (0.1% Deoxycholic Acid, sodium salt, 500mM NaCl, 50mM HEPES, pH 7.5, 1mM EDTA, 1% TritonX-100), LiCL/Detergent Wash Buffer (0.5% Deoxycholic Acid, sodium salt, 250mM LiCl, 1mM EDTA, 0.5% NP-40, 10mM TrisCl pH 8.0), and TE buffer (1mM EDTA, 10mM TrisCl pH 8.0). Chromatin was eluted in TE with 1% SDS at 65°C, and cross-links in both IP and input samples were reversed by overnight incubation at 65°C. Samples were treated with RNase A and proteinase K, and DNA was purified using the Zymo ChIP DNA Clean & Concentrator kit (#D5205). Library preparation was performed using the NEBNext Ultra II DNA Library Prep Kit (NEB, #E7645L/#E7600S). Library quality and concentration was assessed by Qubit and TapeStation prior to sequencing.

### ChIP-Seq Analysis

Libraries were sequenced through Novogene on Illumina NovaSeq X Plus (200G) with 150 base pair paired-end reads. Sequences were aligned to human reference hg38 using BWA (0.7.18).^128^ Samtools (1.21)^139^ was used to remove the PCR duplicates (rmdup) and the reads with a mapping quality score of less than 10 from the aligned reads. Bigwig files of the data generated with deeptools^129^ (v3.5.6, --binSize 10 --effectiveGenomeSize 2913022398 --ignoreForNormalization chrX -extendReads 150) and visualized using pyGenomeTracks. For normalization of the data, each number of the filtered reads was divided by the lowest number of the filtered reads in the same set of experiments, generating a downsampling factor for each sample. Normalized BAM files were generated using samtools view -s with the above downsampling factors and further converted to normalized BAM files using bamCoverage–binSize 10– extendReads 150. Peaks were called by MACS3 where q < 0.05.^131^ Heatmaps were generated from read depth normalized bigwig files using deeptools ComputeMatrix and visualized with plotHeatmap. Motif analysis was done using HOMER’s (5.1) findMotifsGenome command and known motifs were plotted.

#### Polar metabolomics

Cells were plated at 175,000 cells/ml in tissue culture treated, poly-L-lysine-coated 6 cm dishes in DMEM supplemented with 10% dialyzed FBS (three technical replicates were prepared for metabolomics analysis and two technical replicates were prepared for protein quantification for normalization purposes). The following day, sample dishes were placed on wet ice, media was completely aspirated, and cells were washed once with cold PBS. PBS was fully aspirated and plates were transferred to dry ice. Cold 80% methanol stored at −80 °C was added to each dish (1.5 ml per dish). Cells were then scraped on wet ice, and samples were transferred to 2 ml microcentrifuge tubes and vortexed briefly. Samples were centrifuged at 20,000 x g for 5 minutes and immediately dried under nitrogen gas. Dried samples were stored at −80 °C. For LC/MS analysis, dried samples were resuspended in 20 μL of mass spectrometry-grade water, centrifuged at 10,000 x g at 4 °C for 3 minutes, and supernatants were transferred to mass spectrometry compatible vials. Samples were analyzed by the BIDMC Mass Spectrometry Core using a 6500 QTRAP instrument. *Targeted LC/MS metabolomics*.

Samples (5-7 μL) were injected and analyzed using a hybrid 6500 QTRAP triple quadrupole mass pectrometer (AB/SCIEX) coupled to a Prominence UFLC HPLC system (Shimadzu) via selected reaction monitoring (SRM) of a total of 300 endogenous water-soluble metabolites for steady-state analyses of samples. Some metabolites were targeted in both positive and negative ion mode for a total of 311 SRM transitions using positive/negative ion polarity switching. ESI voltage was +4950V in positive ion mode and –4500V in negative ion mode. The dwell time was 3 ms per SRM transition and the total cycle time was 1.55 seconds. Approximately 9-12 data points were acquired per detected metabolite. Samples were delivered to the mass spectrometer via hydrophilic interaction chromatography (HILIC) using a 4.6 mm i.d. x 10 cm Amide XBridge column (Waters) at 400 μL/min. Gradients were run starting from 85% buffer B (HPLC grade acetonitrile) to 42% B from 0-5 minutes; 42% B to 0% B from 5-16 minutes; 0% B was held from 16-24 minutes; 0% B to 85% B from 24-25 minutes; 85% B was held for 7 minutes to re-equilibrate the column. Buffer A was comprised of 20 mM ammonium hydroxide/20 mM ammonium acetate (pH=9.0) in 95:5 water:acetonitrile. Peak areas from the total ion current for each metabolite SRM transition were integrated using MultiQuant v3.0.2 software (AB/SCIEX) *Metabolomics data analysis*.

Metabolomics data were filtered to remove metabolites lowly detected or not detected in at least two technical replicates. Metabolite abundances were normalized to protein content measured using the Bio Rad DC protein assay from parallel harvested samples. Protein normalized metabolite values were used to generate heatmaps in MetaboAnalyst 6.0 using log10 transformation and Pareto scaling.

#### Immunofluorescence

Cells were seeded (50,000-100,000 cells/well) on poly-L-lysine (PLL) coated glass coverslips in a 6 well dish and allowed to adhere overnight under standard culture conditions. Corresponding treatments were performed the next day (e.g. siRNA transfection for 48 hours, using the previously described method), before coverslips were fixed with 4% paraformaldehyde (Millipore-Sigma, 1004965000) for 10 minutes at room temperature. Fixed cells were permeabilized with 0.2% Triton X-100 in OBS for 10 minutes and then blocked with 0.5% bovine serum albumin (BSA) for 1 hour to reduce non-specific binding. Primary antibodies (BRD2: Cell Signaling (5848), BRD2: Abcam (Ab245436), or BRD2: Fortis Life Sciences (A700-008) were diluted 1:100 in blocking solution and applied to coverslips overnight at 4 °C. Coverslips were washed three times with PBS and incubated with fluorescently labeled secondary antibodies (ThermoFisher Scientific, A-11037) for 1 hour at room temperature in the dark.

Coverslips were washed for three times with PBS and mounted onto glass slides using Vectashield mounting media containing DAPI (Vector Laboratories, H-1200-10). Fluorescent images were acquired using a Keyence BZ-X Series miroscoep equipped with a 20x, 40x, or 60x oil immersion objective lens, with exposure time held constant across samples. All images were processed and analyzed in ImageJ/Fiji.

#### Confocal microscopy

BRD2 knock-in mutant cells (Control, S37A and S37D) were seeded onto sterile glass coverslips that were pretreated with Poly-L-Lysine solution (PLL; Sigma-Aldrich, P4707-50ML) for 5 minutes. Cells were seeded in 6-plates with one coverslip/well at a density of 150,000 cells/well and allowed to adhere overnight. Cells were then fixed with 4% paraformaldehyde (PFA; ThermoFisher Scientific, J61899.AK) in PBS for 10 minutes at room temperature. After fixation, cells were washed three times with PBS and permeabilized with 0.2% Triton X-100 (ThermoFisher Scientific, BP151-100) in PBS for 15 minutes on a nutator at room temperature. Coverslips were washed three times with PBS and blocked with 0.5% bovine serum albumin (BSA; Gold Biotechnology, A-420-100) in PBS for one hour on a nutator at room temperature. Coverslips were then incubated overnight on a rocker at 4°C with BRD2 primary antibody (Abcam, Ab245436) diluted in blocking buffer at 1:100. Following primary antibody incubations, coverslips were washed three times in PBS and incubated with a fluorophore-conjugated secondary antibody diluted 1:1000 in blocking buffer (Thermofisher Scientific, A-11037) for one hour at room temperature in the dark. Coverslips were then mounted on slides using 10 µL of DAPI Fluoromount-G (SouthernBiotech, 0100-20). Confocal images were acquired using a Zeiss LSM 880 confocal microscope equipped with a C-Aprochromat 40x/1.2 NA objective lens. All images were taken in the presence of Immersol W (water) immersion oil corresponding to 40x lens, and image settings were kept constant across samples. Images were viewed and processed equivalently using ZEN-Lite software, and scale bars were added using ImageJ/Fiji.

#### Digital Twin Model and In Silico Clinical Trials

##### ODEs of the BETi-PI3Ki TNBC progression model

Here, we derive the ODEs associated with the mechanistic BETi-PI3Ki TNBC progression model introduced in the main paper. The model describes the response of TNBC cells to BET and PI3K inhibition by integrating cell-death dynamics with MYC regulation, chromatin modification dynamics, and BET and PI3K/AKT signaling. The cellular population is represented by a single viable TNBC state and a dying-cell state. Let *Y*_1_(*t*) denote the percentage of dying cells. Because viable and dying cells constitute the total cell population in the model, the percentage of viable TNBC cells can be written as *V*(*t*) = 100 − *Y*_1_(*t*). The transition from the viable state to cell death is regulated by MYC, whose concentration is denoted by *Y*_2_(*t*). Specifically, reduced MYC levels promote the transition toward cell death.^56, 140–142^

To mechanistically represent the epigenetic regulation of MYC, we incorporated a chromatin modification circuit previously introduced,^90^ which was adapted from previous models of chromatin dynamics.^91–93^ For the MYC gene, we consider a nucleosome with DNA wrapped around it as the basic modifiable unit and assume the total number of nucleosomes associated with the gene, denoted by D_tot_, to be constant. Each nucleosome can carry either a repressive (H3K27me3) or an activating (H3K4me3/ac) histone modification. In terms of mechanisms, the circuit includes the competing establishment and maintenance of activating and repressive histone modifications through autocatalytic processes. To reduce the complexity of the model, each nucleosome is assumed to be modified with either an activating or a repressive modification, with no possibility of being unmodified. Under this assumption, loss of one chromatin modification coincides with the establishment of the opposing one, allowing activation and repression to be described through a single state variable while preserving the key mechanisms of chromatin regulation.^90^

We define *Y*_3_(*t*) as the number of nucleosomes with activating histone modifications associated with the MYC gene, while *R_M_*(*t*) = D_tot_ − *Y*_3_(*t*) represents the number of nucleosomes with repressive histone modifications. As in our previous chromatin models,^90, 91^ we assume D_tot_ to be sufficiently large so that these quantities can be treated as continuous variables. In our analyses, we set D_tot_ = 50.^90, 91^

Finally, let *Y*_4_(*t*) and *Y*_5_(*t*) represent the concentrations of functionally active BET and AKT proteins, respectively, and let *I_B_*(*t*) and *I_P_*(*t*) denote the concentrations of the BETi and PI3Ki, respectively. These inhibitor concentrations are constant in the *in vitro* simulations and are determined by the corresponding pharmacokinetic models in the *in silico* clinical trial simulations (see sections below).

Let us now define the parameters of the ODE model:

- *α_M_* (µM/(h ⋅ nucleosome)) denotes the basal gene expression rate of MYC from the active chromatin state.
- *α_PM_* (µM/(h ⋅ nucleosome)) denotes the PI3K/AKT-dependent contribution to MYC production.
- *δ_M_* (h^−1^) denotes the basal decay rate of MYC.
- *δ*_1_ (h^−1^) denotes the maximal rate of treatment-induced death. Additionally, we multiply *δ*_1_ by an exponentially decaying factor *e*^−*mt*^ to model the potential loss of treatment efficacy over time, with *m* > 0 (h^−1^) representing the rate of this decline.
- *δ_d_* (h^−1^) denotes the dying cell clearance rate
- *β_M_* (h^−1^) denotes the basal rate of chromatin activation for the MYC gene.
- 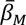 (1/(h ⋅ nucleosome)) denotes the strength of the BET-dependent autocatalytic process promoting establishment and maintenance of the active MYC chromatin state.
- *ε_M_* (h^−1^) denotes the basal rate of chromatin repression at the MYC gene.
- 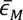(1/(h ⋅ nucleosome)) denotes the strength of the autocatalytic process through which the repressive chromatin state promotes its own establishment.
- *α_B_* (µM/h) denotes the basal production rate of functionally active BET protein.
- *δ_B_* (h^−1^) denotes the maximal BETi-induced reduction rate of functionally active BET protein.
- *α_P_* (µM/h) denotes the basal production rate of AKT.
- *δ_P_* (h^−1^) denotes the maximal PI3Ki-induced decay rate of AKT.
- *α_M_*_→*P*_ (µM/h) denotes the MYC-dependent contribution to PI3K/AKT signaling activity.
- 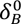 and 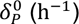 denote the basal decay rate of functionally active BET protein and AKT protein.

The ODEs governing the BETi-PI3Ki TNBC progression model can then be written as follows:

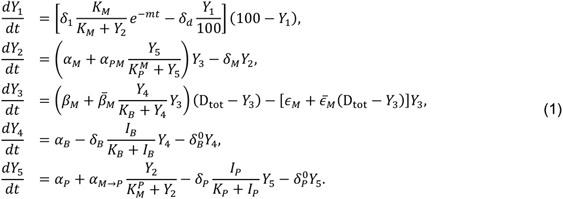

#### Derivation of the BETi-PI3Ki TNBC progression model (SI_PD_model_ODEs)

We first derive the ODE governing the cellular-state variable *Y*_1_(*t*). Let *V̄* (*t*) and *Ȳ*_1_(*t*) denote the absolute numbers of viable and dying TNBC cells, respectively. The cellular-state transition model can be represented as

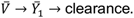

in which the transition from the viable state to cell death is negatively regulated by MYC, and thus its rate can be modeled 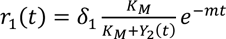 and the dying-cell clearance rate is denoted by *δ_d_*.
The corresponding equations for the absolute numbers of viable and dying cells are

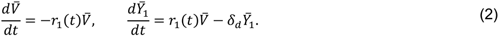

Now, let the total number of cells represented in the system be *C*(*t*) = *V̄* (*t*) + *Ȳ*_1_(*t*), whose dynamics is governed by

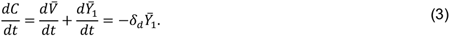

The percentages of *V̄* (*t*) and *Ȳ*_1_(*t*) can then be defined as

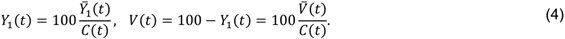

Applying the quotient rule for derivatives to the formula of *Y*_1_(*t*) in (4), we obtain

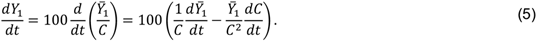

Substituting Eqs. (2) and (3) in Eq. (5), we obtain

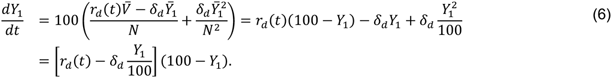

Finally, substituting the expression for *r*_1_(*t*) in Eq. (6), we obtain

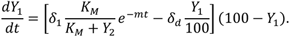

that is, the first equation of the ODE model (1).

We next derive the ODEs governing the molecular species and chromatin-state variables. Here, we define the concentration of functionally active BET protein as *Y*_4_(*t*). Its dynamics are determined by a constant production term *α_B_*, a basal decay with rate 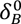, and the effect of BET inhibitors, which bind BET bromodomains and then reduce the amount of functionally active BET protein.^37^ We model this effect through a Hill function^143^ that saturates with the concentration of BETi, *I_B_*. The resulting ODE is

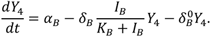

Concerning AKT, its concentration is represented by *Y*_5_(*t*). Its dynamics include basal production, regulation by MYC, basal decay, and the effect of PI3K inhibition. The experimentally observed interaction between BET inhibition and PI3K/AKT pathway reactivation (**Fig. 3D**) is incorporated through MYC-dependent regulation of AKT. Specifically, the MYC-dependent contribution is modeled through the increasing Hill function 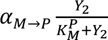. Concerning the effect of PI3Ki, *I_p_*, it inhibits PI3K, thereby reducing downstream AKT activation. This effect is captured by the saturating function 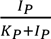. The resulting ODE can then be written as

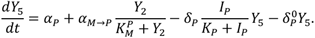

Let us now derive the ODEs associated with MYC. Its concentration is represented by *Y*_2_(*t*). For its production, we consider a basal contribution, with rate *α_M_*, together with an AKT-dependent contribution. The latter is represented by an increasing Hill function of AKT concentration, that is 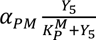. Combining these contributions with basal MYC degradation, we obtain

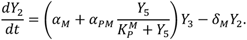

Concerning the chromatin modification dynamics, transition from the repressive to the activating state occurs through two mechanisms: a basal activation process, with rate *β_M_*, and a autocatalytic process promoted by existing activating modifications.^91–93^ In this model, this process is modulated by BET, which recognizes acetylated histones and facilitate recruitment of transcriptional machinery at active chromatin regions.^37–40, 56^ Combining these effects, the effective activation rate can be written as

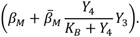

Conversely, transition from the activating to the repressive state occurs through a basal process with rate *ɛ_M_* and an autocatalytic process mediated by existing repressive chromatin. The effective repression rate is therefore 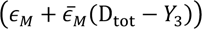). Since these rates act on nucleosomes carrying the opposite modification, the activation and repression rates are multiplied by D_tot_ − *Y*_3_ and *Y*_3_, respectively. The complete equation governing the MYC chromatin state is therefore

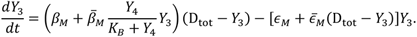

#### PK model of GDC-0077

To model the pharmacokinetics (PK) of inavolisib (GDC-0077), we relied on the human pharmacokinetic analysis reported in [Salphati2024], together with the clinical pharmacology information reported in the FDA prescribing information.^96^ Salphati et al.^144^ developed a physiologically based PK model to predict the human plasma concentration-time profile of inavolisib and subsequently fitted the predicted profile using a one-compartment model with first-order oral absorption.^144^ The resulting model provided estimates of the absorption rate constant, elimination rate constant, and apparent volume of distribution and showed good agreement with plasma concentrations subsequently measured in patients [Salphati2024]. We therefore adopted the same one-compartment structure.

In order to introduce the ODE model, let *A*_GI_(*t*) denote the amount of inavolisib in the gastrointestinal compartment and *C_p_*(*t*) its plasma concentration. The PK model can be written as

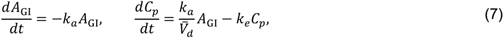

where *k_a_* denotes the first-order absorption rate constant, *k_e_* the first-order elimination rate constant, and *V̄_d_* = *V_d_*/*F* the apparent volume of distribution, with *F* representing the bioavailability. The parameter values were taken directly from [Salphati2024], namely *k_a_* = 0.73h^−1^, *k_e_* = 0.046 h^−1^, and *V_d_*/*F* = 245 L. These parameters correspond to an apparent oral clearance of *CL*/*F* = *k_e_*(*V_d_*/*F*) = 11.27 L/h.

Because the model is parameterized in terms of the apparent volume of distribution, i.e., *V̄_d_* = *V_d_*/*F*, oral bioavailability is implicitly incorporated and thus administered oral doses were introduced directly without applying an additional bioavailability correction. At each dosing event, the gastrointestinal amount was updated according to

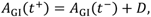

where *D* denotes the administered dose.

To account for inter-individual pharmacokinetic variability, patient-specific values were sampled from log-normal distributions.^145^ Specifically, each parameter *P_i_* was generated according to

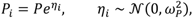

where *P* denotes the nominal population value and *ω_P_* was calculated from the corresponding coefficient of variation, CV*_P_*, as

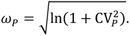

The CVs of 26% for the apparent volume of distribution and 29% for clearance were taken from inavolisib prescribing information as defined by the FDA.^96^ For each virtual patient, the elimination rate constant was then calculated as

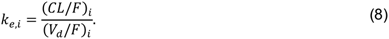

Because inter-individual variability in the absorption rate constant was not reported in the available clinical sources, we introduced a modest assumed variability of 10% around the population value of *k_a_* = 0.73 h^−1^.

For each virtual patient, the resulting PK model was simulated to obtain the corresponding plasma concentration profile of inavolisib over time.

#### PK model of ZEN-3694

To build the PK model of ZEN-3694, we relied on the clinical pharmacokinetic data reported in previous studies.^145, 146^ Aggarwal et al.^145^ characterized ZEN-3694 PK and its active metabolite ZEN-3791 across multiple dose levels, while, more recently, Feng et al.^146^ reported separate plasma concentration-time profiles for ZEN-3694 and ZEN-3791 following administration of a single 48 mg oral dose. We used the parent-specific ZEN-3694 profile reported in Feng et al.^146^ to parameterize a one-compartment model with first-order oral absorption and elimination. This model was selected because the observed parent-drug profile showed a clear absorption phase followed by a smooth decline, without evidence from the available data that would justify the introduction of an additional systemic compartment. The PK model can then be written as in (7).

To estimate the model parameters, we used the parent ZEN-3694 concentration-time profile reported in Figure 4 of Feng et al.^146^ following administration of a 48 mg oral dose. From this profile, we estimated a maximum plasma concentration of approximately 487 ng/mL at 2–3 hr and an AUC_O−24_ of approximately 3.48 mg⋅h/L. The apparent oral clearance was estimated from the administered dose and total parent-drug exposure as *CL*/*F* = *D*/AUC_O−∞_ ≈ 13.6 L/h.^147^ The terminal decline of the parent-specific concentration profile was used to estimate an elimination half-life of approximately 3.5 h, corresponding to *k_e_* = ln(2)/*t*_1/2_ ≈ 0.198 h^−1^. From the estimated apparent clearance and elimination rate, the apparent volume of distribution was calculated as *V̄_d_* = *V_d_*/*F* = (*CL*/*F*)/*k_e_* ≈ 68.7 L. Furthermore, based on the observed absorption phase and *T*_max_ of approximately 2–3 h, the absorption rate constant was set to *k_a_* = 1.0 h^−1^. Thus, the nominal PK parameters used in the simulations were *k_a_* = 1.0 h^−1^, *k_e_* = 0.198 h^−1^, and *V̄_d_* = 68.7 L.

Finally, although ZEN-3694 has an active metabolite, ZEN-3791,^145, 146^ this was not explicitly included in this PK model because the available clinical data were insufficient to reliably parameterize a mechanistic parent–metabolite model.

As for the PK model of GDC-0077, oral bioavailability was directly incorporated into the model through the apparent volume of distribution. Thus, at each dosing event, the gastrointestinal amount was updated according to

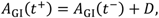

where *D* denotes the administered dose.

To account for inter-individual pharmacokinetic variability, we used the same approach used for the GDC-0077 PK model. Given that parameter-level estimates of inter-individual pharmacokinetic variability were not available from the identified clinical studies, for this model modest variability was considered to generate the virtual patient population and evaluate the robustness of the treatment optimization. Specifically, the apparent clearance and apparent volume of distribution were assigned CVs of 20%, while the absorption rate constant was assigned a CV of 30%. For each virtual patient, the elimination rate constant was then calculated as in (8) and the resulting PK model was simulated to obtain the corresponding plasma concentration profile of ZEN-3694 over time.

#### *In silico* clinical trials

The parameter estimation for the BETi-PI3Ki TNBC progression model, *in silico* clinical trials, and treatment schedule optimization were performed as described previously.^90^

